# Quantification of surface tension effects and nucleation-and-growth rates during self-assembly of biological condensates

**DOI:** 10.1101/2022.11.23.517626

**Authors:** Zsuzsa Sárkány, Fernando Rocha, Anna Bratek-Skicki, Peter Tompa, Sandra Macedo-Ribeiro, Pedro M. Martins

## Abstract

Liquid-solid and liquid-liquid phase separation (PS) drives the formation of functional and disease-associated biological assemblies. Principles of phase equilibrium are here employed to derive a general kinetic solution that predicts the evolution of the mass and size of biological assemblies. Thermodynamically, protein PS is determined by two measurable concentration limits: the saturation concentration and the critical solubility. Due to surface tension effects, the critical solubility can be higher than the saturation concentration for small, curved nuclei. Kinetically, PS is characterized by the primary nucleation rate constant and a combined rate constant accounting for growth and secondary nucleation. It is demonstrated that the formation of a few number of large condensates is possible without active mechanisms of size control and in the absence of coalescence phenomena. Our exact analytical solution can be used to interrogate how the elementary steps of PS are affected by candidate drugs.

## Introduction

Multiple lines of evidence indicate that liquid-liquid phase separation (LLPS) of proteins has an important role in the spatiotemporal regulation of cell functioning and disease development [1]. These findings motivate a renewed interest in the detailed description of the nucleation-and-growth mechanisms by which new protein phases are generated. Given the tinctorial properties of amyloid fibrils, a large number of kinetic curves have been recorded to characterize protein aggregation *in vitro* using thioflavin-T and other specific dyes [2]. These studies are important to understand the nucleation-and-growth of amyloid fibrils and other insoluble deposits associated with neurodegenerative diseases. Biophysical models such as those proposed by Crespo et al. [3,4] and by Finke and Watzky [5,6], can describe, at least qualitatively, the hyperbolic, S-shaped and sigmoidal progress curves usually reported for peptides and proteins such as amyloid-β peptide (Aβ), α-synuclein, tau, prion, transthyretin (TTR), insulin, ataxin-3, β_2_-microglobulin, huntingtin, etc. In the case of sigmoidal curves, the nucleation-and-growth mechanism is further validated by scaling laws of half-life coordinates and maximal aggregation rate vs. protein concentration [7,8], and by measurements of particle size over time [9]. Although less frequently observed, hyperbolic-shaped progress curves characterize the amyloid aggregation of TTR [10], as well as the aggregation of Aβ at high peptide concentrations [11] or in the presence of amyloid inhibitors [12].

Hyperbolic, S-shaped and lag-phased curves are also expected by the nucleation-and-growth model proposed by Zwicker et al. [13] intended to describe the sigmoidal growth of centrosomes as an LLPS process controlled by an enzymatic activity of two centrioles within the centrosome matrix. Other membraneless organelles, like nuclear bodies and cytoplasmic granules undergo coacervation through the recruitment of protein and RNA molecules in supersaturated solutions, giving rise to a power-law dependence of the average droplet size 〈*R*〉 on time *t*. If a fixed number of growing droplets is assumed, the time exponent *n* takes on values of ~1/2 in purely diffusional regimes and ~1 in kinetic regimes determined by surface attachment [14]. The *n* = 1/2 exponent is also predicted by the theory of diffusion-limited precipitation, which, in addition, rationalizes the application of the Johnson–Mehl–Avrami–Kolmogorov (JMAK) equation to the early stages of LLPS progress curves [14]. Both the JMAK- and logistic-type equations predict that the supersaturated phase becomes fully depleted of protein for long reaction times or, equivalently, that the cellular compartment becomes completely filled with the droplet phase [4,14]. In order to numerically fit experimental data, these equations can be modified by the inclusion of corrected asymptotic limits for *t* → ∞ [14,15], whereas a power-law relationship between 〈*R*〉(*t*) and the reaction conversion are admissible if the number of growing particles is approximately constant [6]. After the nucleation-and-growth period has passed, the initially supersaturated phase becomes saturated and the volume fraction of the new phase increases no more. From this point onwards, the average size of dense droplets in saturated solutions can continue to increase due to Ostwald ripening and/or coalescence processes in which smaller droplets convert to fewer and larger droplets driven by surface tension effects. The coalescence period that follows nucleation-and-growth is well characterized by a dynamic scaling exponent *n* of 1/3 [14].

The nucleation-and-growth of a few isolated particles without the continual formation of new nuclei is a challenge to the existing kinetic models with important implications for our understanding of pathogenic protein aggregation and cell organization. The time and extension of nucleation can be regulated by enzymatic activity [13], parallel oligomerization reactions [8] and ATP-driven reactions [14] that indirectly affect the supersaturation level. New techniques and methods for studying LLPS show, however, that the formation of a reduced number of non-coalesced condensates is possible in the absence of external sources of energy or matter [15–20]. To help understand these intriguing observations, we start by including surface tension effects (STEs) on established nucleation-and-growth models; the resulting closed-form kinetic equations are then tested against LLPS progress curves and are used to predict the variation with time of droplet size distributions. Finally, we discuss fundamental and biological implications arising from STEs-affected assembly and summarize the steps that should be taken to quantify STEs and nucleation-and-growth rates.

### Thermodynamics and kinetics of nucleation-and-growth

The formation of liquid protein phases follows thermodynamic and kinetic principles that are common to other phase-transition phenomena. For example, the theoretical free energy of the mixing of binary polymer-solvent solutions can be used to estimate the equilibrium concentrations of the protein-depleted (or *light*) and protein-enriched (or *dense*) phases formed upon LLPS [21,22]. In temperature vs. composition phase diagrams, these concentrations are generally expressed in terms of the protein volume fractions *φ_L_* and *φ_D_*, and define the so-called *coexistence curve* or binodal (Fig. 1a). For attractive interactions between molecules, there is a critical temperature *T_c_* above which phase separation does not occur, regardless of the concentration [23,24]. LLPS provoked by increasing temperatures can, however, occur in the case of predominantly hydrophobic interactions [25]. In liquid-solid phase diagrams, the equilibrium compositions of the light (liquid) and dense (solid) phases are given by the *solubility curve* (or liquidus) and by the solidus line, correspondingly (Fig. 1b). The volume fraction of the protein in the solid phase does not change significantly with temperature and, in the case of protein crystals, can be lower than 0.5. Liquid-liquid immiscibility can be metastable in relation to the solid phase, in which case LLPS is a precursor step of the more favourable liquid-solid transition [24,26,27]. According to the phase rule of thermodynamics, the number of possible microstates due to demixing increases with the number of non-reactive components present in the mixture [28]. Consequently, the complexity of the phase diagrams increases dramatically by considering the occurrence of interaction partners and metabolites, but also by including the effect of pH, ionic strength and additives [29], possible eutectic behaviours [30], the spontaneous formation of the new phase by spinodal decomposition [28], etc. In what follows, we will address the regions of protein phase diagrams where the formation of stable particles (protein crystals, aggregates or droplets) takes place by a nucleation-and-growth mechanism.

**Figure 1.**
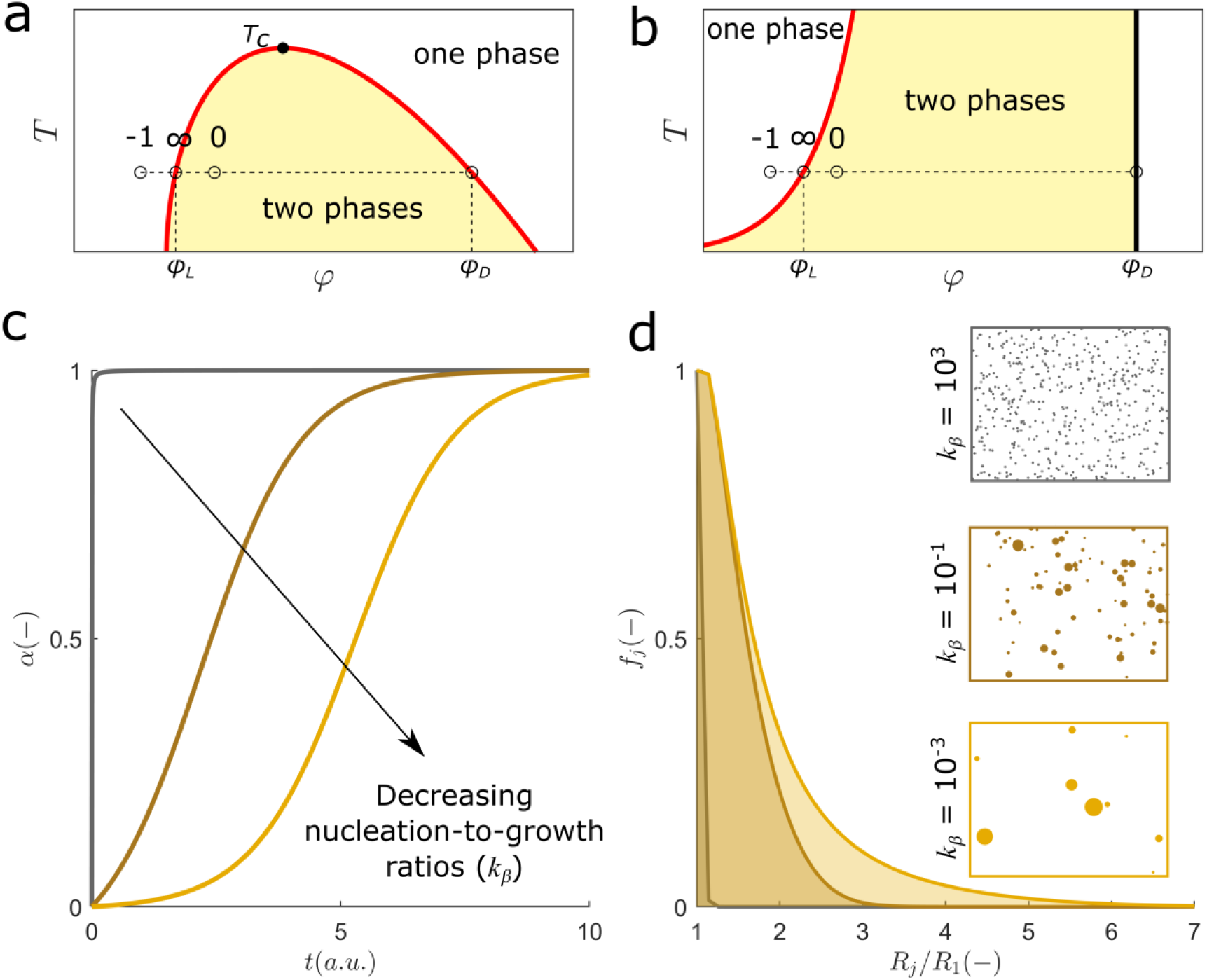
Thermodynamics and kinetics of nucleation-and-growth. (a and b) Schematic representation of phase diagrams of binary protein-water mixtures representing temperature (*T*) vs. volume fraction of the protein (*φ*). Points along the dashed lines represent undersaturated (−1), supersaturated (0) and saturated conditions (∞) at a fixed temperature. (a) liquid-liquid equilibrium: the volume fraction of the protein in the light (*φ_L_*) and in the dense (*φ_D_*) phase are given by the coexistence curve (solid line) that shows a maximum at the critical temperature *T_c_*. (b) liquid-solid equilibrium: the values of *φ_L_* and *φ_D_* are given by solubility curve (red line) and by the solidus line (black line), respectively. (c) Theoretical progress curves for nucleation-to-growth ratios of (from left to right) *k_β_* = 10^3^, 10^−1^ and 10^−3^ (*SI Additional Methods*). (d) Normalized frequency (*f_j_*) of particles with size *R_j_* calculated for long reaction time (*t* → ∞) using the same parameters and colour code as in (c); *R*_1_ is the size of primary nuclei. Cartoon boxes: the final population of particles is schematically represented as a distribution of circles. No secondary events such as secondary nucleation, breakage or coalescence are considered in (c) and (d).

Consider the change from the one-phase to the two-phase regime of the phase diagram by means of the isothermal increase of protein concentration from *φ*_−1_ < *φ_L_* to *φ*_0_ > *φ_L_*. If the temperature is kept constant, the system will naturally relax towards equilibrium through the nucleation and subsequent growth of a dense phase that has constant composition (*φ_D_*) and decreases the protein volume fraction in the light phase *φ* until saturated conditions are reached (*φ*_∞_ = *φ_L_*). The individual rates of the nucleation and growth steps determine how fast the change from *φ*_0_ to *φ*_∞_ is, whereas the ratio of nucleation-to-growth rates (*k_β_*) determines the shape of phase separation progress curves. In the absence of other events, increasing values of *k_β_* lead to a transition from hyperbolic, to S-shaped to lag-phased progress curves (Fig. 1c) [3]. In concentration units, the amount of protein separated into the new phase is *M*(*t*) = *c*_0_ – *c*(*t*), where *c*_0_ and *c*(*t*) are the initial and instantaneous protein concentrations; *M*(*t*) increases with time until reaching the final value of *M*_∞_ = *c*_0_ – *c*_∞_. In the phase diagram, c_0_ and *c∞* correspond to volume fractions *φ*_0_ and *φ*_∞_ = *φ_L_*, respectively. The instantaneous reaction conversion is *α*(*t*) = *M*(*t*)/*M*_∞_. For large particles to be formed, primary nucleation should be much slower than the growth step (*k_β_* ≪ 0.01) so that early nucleated species can continue to grow during the whole duration of lag-phased progress curves. Because of the long period required for phase separation, large particles end up coexisting with recently formed ones that have much smaller sizes (Fig. 1d). If, instead, primary nucleation is fast compared with the growth step (high *k_β_*), numerous small condensates will be produced until phase equilibrium is ultimately achieved (Fig. 1d). In this context, the sharp formation of a few number of large particles is an apparent paradox, whose solution is here sought using principles of phase equilibrium at curved interfaces.

### Surface Tension Effects (STEs) on Nucleation-and-Growth

We investigated how STEs might affect the early steps of phase separation. Besides driving Ostwald ripening [31] and droplet coalescence in later stages of LLPS [32], surface tension is also expected to influence the initial assembly of critically-sized nuclei [33,34]. In fact, the condition of mechanical equilibrium between phases establishes that the difference in fluid pressure inside and outside a curved surface increases as the radius of curvature decreases. For a spherical surface of radius *r*, this positive difference is given by the Young-Laplace equation Δ*p* = 2*γ*/*r*, where *γ* is the surface tension. Consequently, the formation of primary nuclei with a radius of curvature *r* ≈ *R_c_* is driven by a difference in chemical potential Δ*μ_c_* that is lower than the one corresponding to a flat interface (*r* = ∞). By combining the Young-Laplace equation with the condition of chemical equilibrium between phases (Δ*μ* = 0), an equation of the Ostwald-Freundlich type is obtained [35], relating the critical protein solubility *c_c_* with saturation concentration expected for flat interfaces (*c*_∞_):

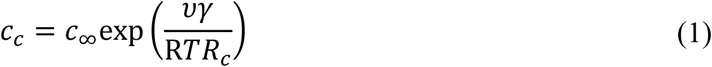

where *ν* is the molar volume of the protein and R is the universal gas constant. This equation estimates the protein solubility enhancement elicited by critically-sized nuclei relative to larger particles characterized by *r* ≈ ∞. Taking as reference the values of *γ* = 0.5 mJ/m^2^ estimated for lysozyme crystals [26], and of *ν* = 0.01 m^3^/mol, a 2% increase in protein solubility is predicted for values of *R_c_* = 100 nm that are close to the initial sizes of amyloid aggregates and of liquid droplets measured by dynamic light scattering [8,16]. The enhancement of protein solubility will be as large as 21% if an estimate of *R_c_* = 10 nm is used in Eq. 1.

The predicted magnitude of STEs is contingent on fixed parameters such as *γ* or *R_c_*, but is also greatly dependent on particle geometry premises. It is likely, for example, that flatter secondary nuclei generated at the surface of the new phase would be less affected by STEs than sphere-like primary nuclei. On the other hand, even slight enhancements of protein solubility will significantly affect the variation in chemical potential during phase separation, which is generally approximated by the value of supersaturation [3]. Supersaturation can also be expressed in concentration units normalized to the initial conditions (subscript 0):

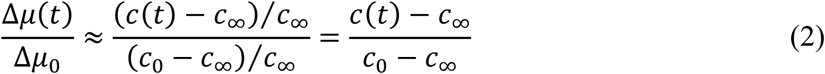

When STEs affect primary nucleation, this typical definition has to be modified taking into account the critical solubility *c_c_* corresponding to the protein concentration in equilibrium with critically-sized particles:

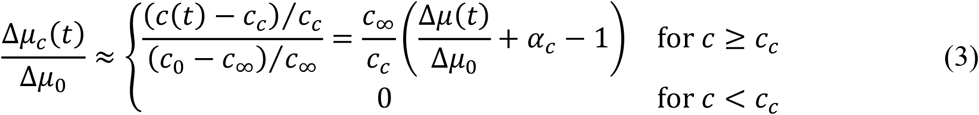

where Δ*μ*(*t*)/Δ*μ*_0_ is given by Eq. 2, and the critical conversion *α_c_* is:

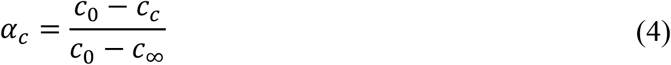

As implied from the above definitions, primary nucleation ceases to occur when the protein concentration decreases below *c_c_* or, equivalently, when the reaction conversion increases above *α_c_*. Thus, STEs can be quantified as the reaction extent during which no primary nucleation is observed: STEs = 1 – *α_c_*.

Nucleated particles are the result of successive monomer addition events until the critical nucleus size is attained and intermolecular forces within the new phase start to prevail over the forces established with the surrounding solution [36,37]. Primary nucleation of macromolecules may also involve substantial structural rearrangements as in the case of amyloid fibril formation [38]. Although primary nucleation is a multistep process, the stochastic assembly of molecules that leads to the formation of a stable nucleus is generally considered rate-limiting owing to the high energy cost required for generating the new interface. In the presence of size-dependent STEs, the energetic cost will be even higher owing to the solubility enhancement caused by small, curved interfaces. It is expected that once this energetic barrier is surpassed, a period of very fast growth will follow catalyzed by the exponential decrease of solubility, and will continue until the radius of curvature is fully relaxed. In such cases, the timescale for primary nucleation is still determined by the initial formation of critically-sized nuclei, but the end-product of the nucleation process consists of particles with size *R*_1_ larger than *R_c_*. As these particles are sufficiently flat and/or large to be insensitive to STEs, their subsequent growth will be driven by the concentration difference (*c*(*t*) – *c*_∞_), until the saturation concentration *c*_∞_ is eventually reached. Therefore, in contrast to what happens for primary nucleation, the driving force for growth can still be defined using Eq. 2.

Departing from the Young-Laplace treatment of curved interfaces, we can conclude that the classical definition of driving force for primary nucleation has to be sometimes corrected to account for the enhancement of protein solubility caused by shape- and size-dependent surface tension effects. This enhancement can be pronounced in the case of small, spherical nuclei but rapidly vanishes as flatter geometries are considered. The existence of a critical solubility value *c_c_* higher than the saturation concentration *c*_∞_ implies that primary nucleation ceases to occur for sub-critical protein concentrations, i.e., only secondary processes take place for protein concentrations < *c_c_*.

### Exact Solutions of the General Nucleation-and-Growth Mechanism

To quantitatively evaluate the impact of STEs, previously established mass balance equations describing the nucleation-and-growth of crystals and amyloid fibrils [4,9] are now modified to include the generalized definition of the driving force for primary nucleation (Eq. 3 and Appendix: *General Nucleation-and-Growth Model*):

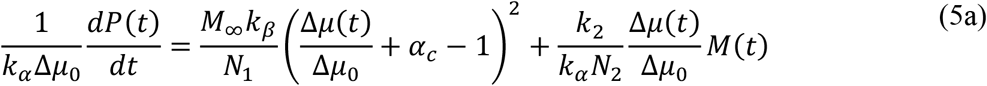

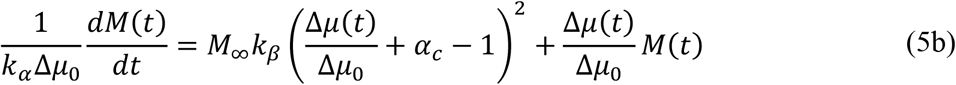

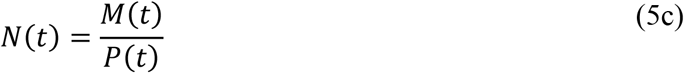

where *P*(*t*) is the number concentration of particles; *N*(*t*) is the mean particle size expressed in terms of number of condensed units; *N*_1_ and *N*_2_ are the number of condensed units constituting the primary- and secondary-nucleation particles, respectively; and *k_α_*, *k_β_* and *k*_2_ are *c*_0_-independent rate constants describing the autocatalytic, primary nucleation and secondary nucleation steps. Negligible secondary nucleation rates (*k*_2_ ≈ 0) will be assumed henceforth as one of our goals is to describe nucleation-and-growth processes producing large condensates, whilst numerous particles with mean size close to *N*_2_ are expected in the presence of increasingly stronger secondary nucleation (Fig. S4) The nucleation- and-growth mechanism underlying the above differential equations is here called “general” because it applies to liquid-solid phase separation, but also to LLPS processes that, in principle, are more prone to STEs provoked by the curvilinear interface of liquid droplets. Despite that, the generalizations and simplifying hypotheses adopted to derive Eqs. 5a and 5b limit the applicability of this mathematical model to cases where: (i) primary nucleation is described by second-order rate equations on supersaturation, (ii) growth and secondary nucleation are autocatalytic processes whose rates increase with the mass concentration of particles and are first-order functions of supersaturation, (iii) *P*(*t*) is not significantly affected by the occurrence of breakage/fragmentation events, nor by coalescence phenomena, and (iv) the volume of the supersaturated solution remains approximately constant [4,9]. After replacing the definitions of Δ*μ*(*t*)/Δ*μ*_0_ = 1 – *α*(*t*) in Eq. 5b, a closed-form *α(t*) expression is obtained upon analytical integration (Fig. 2a) that is dependent on the time-variables *t*_1_, *t_c_* and *τ* (Fig. 2b). The critical instant *t_c_* marks the transition from the nucleation-and-growth to the growth-only regime and coincides with the moment the protein concentration *c*(*t*) equals *c_c_* and the reaction conversion is *α*(*t_c_*) = *α_c_*; the characteristic time constant *τ* determines the timescale for phase separation and reduces to 2(*k_α_*Δ*μ*_0_)^−1^ for *α_c_* = 1; *t*_1_ is the instant of maximum *dα*(*t*)/*dt* slope as given by the split *α*(*t*) expression for *t* ≤ *t_c_*. Note that the products *k_α_*Δ*μ*_0_*t_c_*, *k_α_*Δ*μ*_0_*t*_1_ and *k_α_*Δ*μ*_0_*τ* are functions of two parameters only, *α_c_* and *k_β_*. The representation of the average particle size 〈*R*〉 as a function of *k_α_*Δ*μ*_0_-normalized units of time shows that phase separation becomes slower as STEs get stronger (Fig. 2c). The particle growth curves confirm that the presence of STEs induces the formation of larger particles over more prolonged timescales (Fig. 2d). Similar effects are produced if different nucleation-to-growth ratios are considered (see Figs. 3c and 3d below). Overall, the kinetics of phase separation is determined by the initial supersaturation Δ*μ*_0_, by two kinetic parameters *k_α_* and *k_β_* that are independent of the initial protein concentration *c*_0_, and by the *c*_0_-dependent value of *α_c_*.

**Figure 2.**
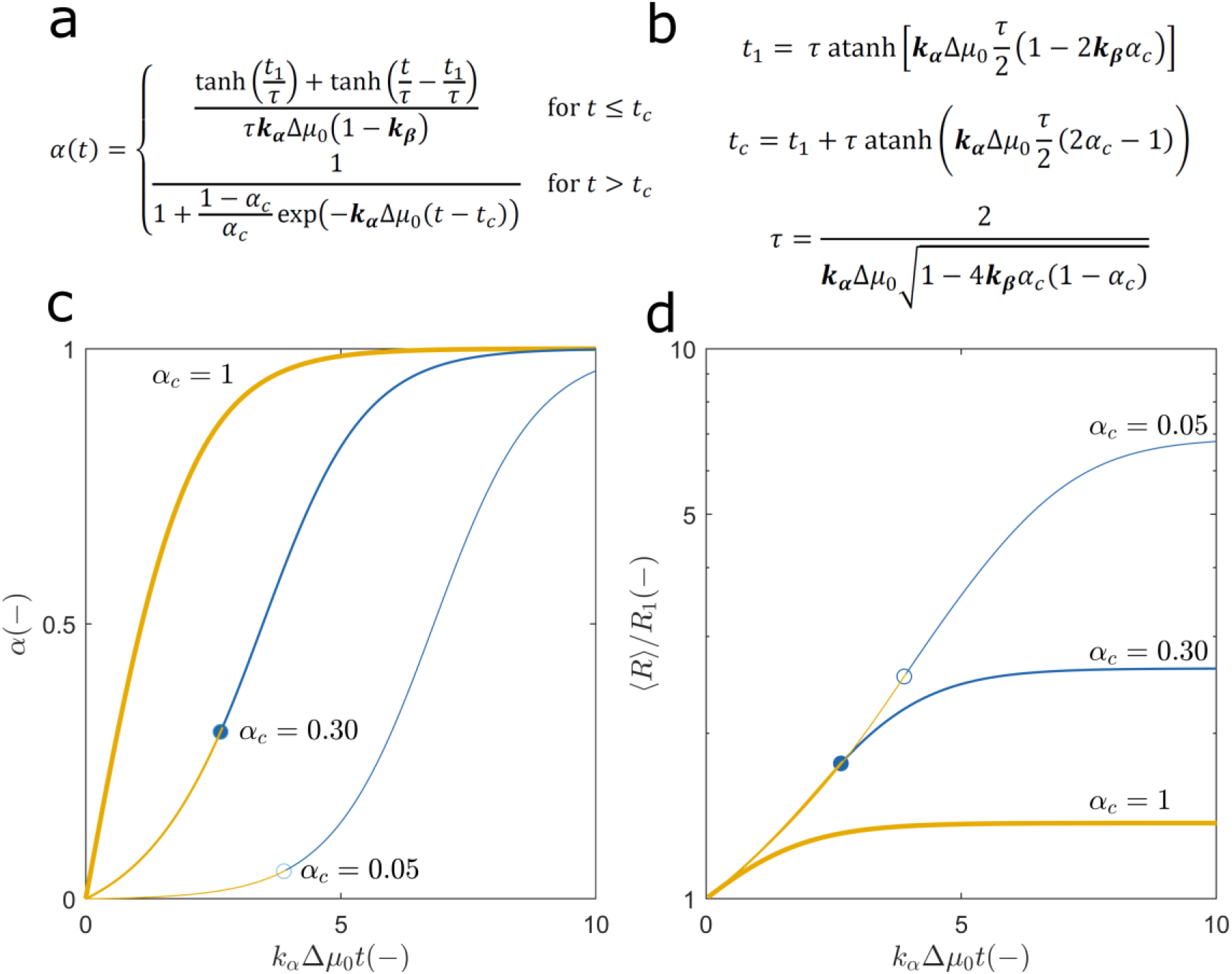
Exact solutions of Eq. 5 describing the nucleation-and-growth of particles in the presence of surface tension effects. (a) Analytical solution of Eq. 5b for the initial condition *α*(0) = 0. (b) Definitions of time-variables *t*_1_, *t_c_* and *τ* in (a) as a function of two *c*_0_-independent kinetic constants, *k_α_* and *k_β_* (in bold). (c) Representation of theoretical progress curves using (from left to right) *α_c_* = 1, 0.30 and 0.05 for a fixed nucleation-to-growth ratio (*k_β_* = 0.5). Lines of different color represent the nucleation-and-growth period (orange) and the growth-only period (blue). Symbols indicate the location of the critical moment *t_c_*. (d) Log-linear plot of the particle growth curves obtained by numeric integration of Eqs. 5a and 5c assuming the same model parameters as in (c). Symbols, line thicknesses and line colors are as in (c).

**Figure 3.**
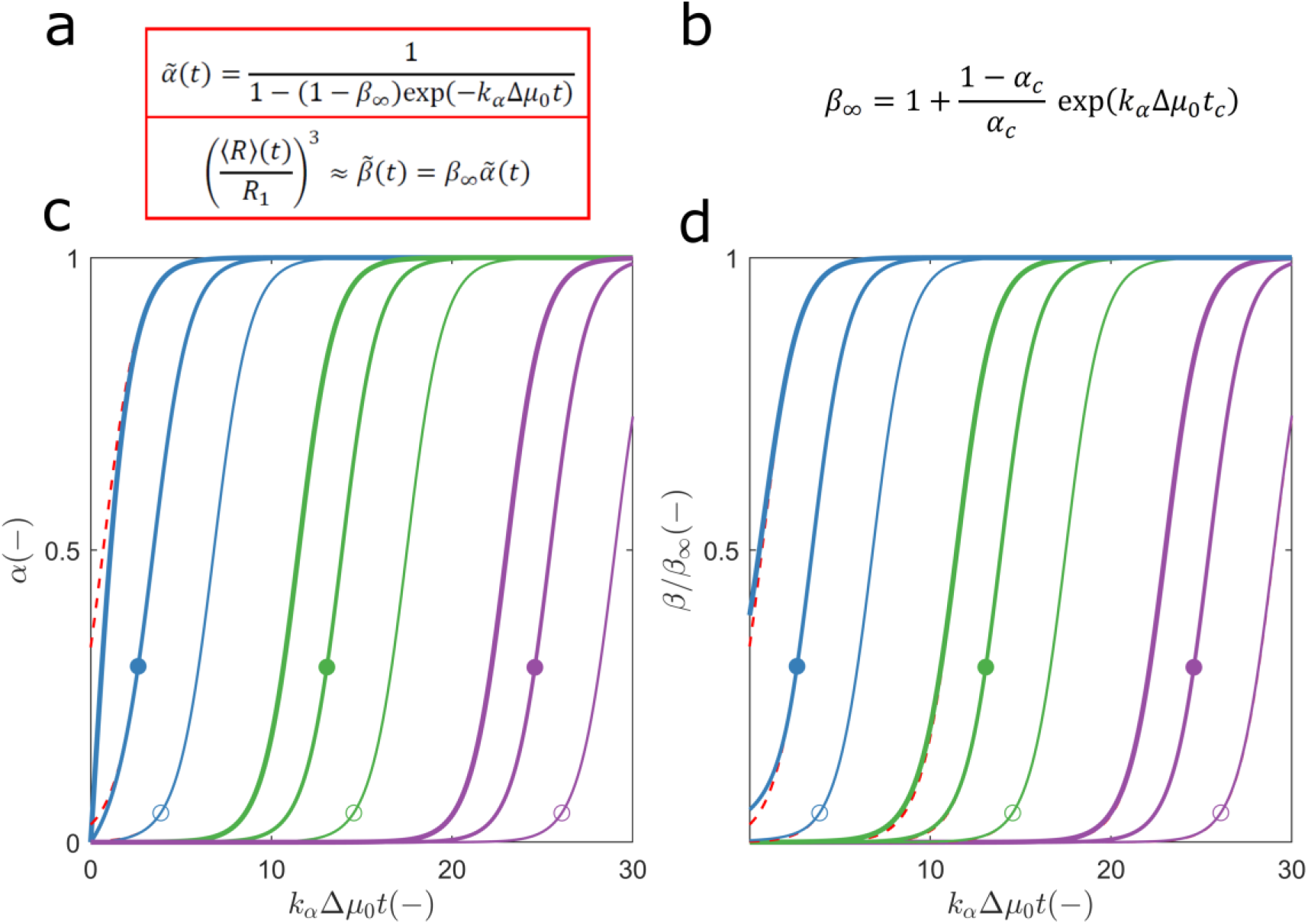
Exact and approximate solutions of Eq. 5. (a) The approximate 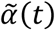 and 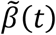 expressions are derived by assuming a constant number of particles. (b) From this definition of the parameter *β*_∞_, it follows that the expression for 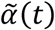 and 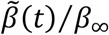 in (a) is equivalent to the exact *α*(*t*) solution derived for *t* > *t_c_* (Fig. 2a). (c) Solid lines: exact *α*(*t*) profiles computed as described in Fig. 2c using *k_β_* values of (different colours from left to right) 0.5, 10^−5^, and 10^−10^, and *α_c_* values of (decreasing order of line thickness) 1, 0.30 and 0.05. Dashed red lines (overlapped when not seen): 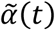 profiles calculated from the definitions in (a) and (b) using the same values of *k_β_* and *α_c_* as in the exact solution. Symbols: location of the critical coordinates *t_c_* and *α_c_*. (d) Comparison between the exact *β*(*t*) = *N*(*t*)/*N*_1_ (solid lines) and approximate 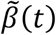 (dashed red lines) solutions calculated using the same model parameters as in (c).

The occurrence of STEs may result in the rapid nucleation and subsequent growth of a reduced number of large particles; it can also explain unconventional effects of protein concentration on the measured *α*(*t*) curves. Before illustrating these possibilities with practical examples, approximate solutions of the *N*(*t*) and 〈*R*(*t*)〉 curves are provided next towards a simple and general methodology for phasese-paration kinetic analysis.

### Particle Growth Curves - Approximate Solution

Unlike the *α*(*t*) solution, the analytical *N*(*t*) solution of Eqs. 5a and 5c is too complex to be used in routine analyses of growth curves as an alternative to the exact numerical solution. An approximate expression is now derived, assuming a constant number of growing particles *P*(*t*) ≈ *P*_∞_, which implies a final mean size of *N*_∞_ = *M*_∞_/*P*_∞_. On this basis, the exact *α*(*t*) equation presented in Fig. 2a for the growth-only period can be extended to the whole nucleation-and-growth period (Fig. 3). By combining *k_α_*Δ*μ*_0_, *k_β_* and *α_c_* in a single variable *β*_∞,_ an approximate 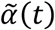 expression is obtained (Figs. 3a and 3b) containing two model parameters (*β*_∞_ and *k_α_*Δ*μ*_0_) that can be numerically fitted to later stages of *α*(*t*) curves (Fig. 3c). Direct replacement of this expression in Eq. 5c gives the equality 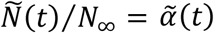, whose limit for *t* → 0 leads to the result 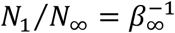. The expression for 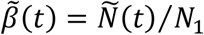 is appropriate for estimating *β*_∞_ and *k_α_*Δ*μ*_0_ from droplet size measurements (Figs. 3a and 3d).

The differences between the exact and approximate solutions are more pronounced during the initial periods of fast nucleation processes and, as expected, are null during growth-only periods, independently of the considered value of *k_β_* (Figs. 3c and 3d). The application of the 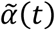 expression to the early stages (*t* < *t_c_*) of hyperbolic progress curves is disqualified by the high errors associated with this approximation (Fig. S5a). On the contrary, the 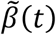 approximation adequately describes the generality of particle growth curves (Fig. S5b). In the worst possible scenario, the 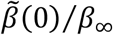 estimates give approximate errors of −8% or +8% for values of *k_β_* ≈ 10^−1^ or *k_β_* ≈ 10, respectively, if STEs are absent (Fig. S5b, inset). In all cases, initial differences between *β*(*t*) and 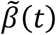 vanish as phase separation progresses (Fig. 3d).

### Kinetic Analysis of LLPS

The new insights provided by the general nucleation-and-growth model are first illustrated using LLPS kinetic data obtained for the low-complexity domain (LCD) of heterogeneous nuclear ribonucleoprotein A2 (hnRNPA2), the TAR DNA-binding protein 43 (TDP-43), the nuclear pore complex protein NUP98 and the Early Responsive to Dehydration 14 (ERD14) stress protein [16]. These aggregation-prone proteins can be kept stable in solution at a carefully selected extreme pH and then undergo LLPS by imposing native pH conditions through the addition of a small amount of concentrated buffer [16]. Particle growth curves measured by dynamic light scattering (DLS) reveal that droplets with initial sizes > 100 nm grow at different rates in the presence and the absence of 150 mM NaCl (Fig. 4). As an alternative to empirical 〈*R*〉 vs. *t* scaling laws, the 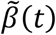 expression in Fig. 3a can be used to fit the measured data considering that 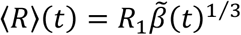 for invariant droplet shapes. Parameters *β*_∞_ and *k_α_*Δ*μ*_0_ are numerically fit to the experimental 〈*R*〉(*t*) curves using the observed initial size as an estimate of 〈*R*_1_〉. When the final size 〈*R*〉(∞) is also known, the value of *β*_∞_ can be estimated in advance from the relationship 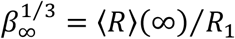. This combined parameter is affected both by *α_c_* and *k_β_* and represents the number of times the droplet volume is ultimately increased (Fig. 3b).

**Figure 4.**
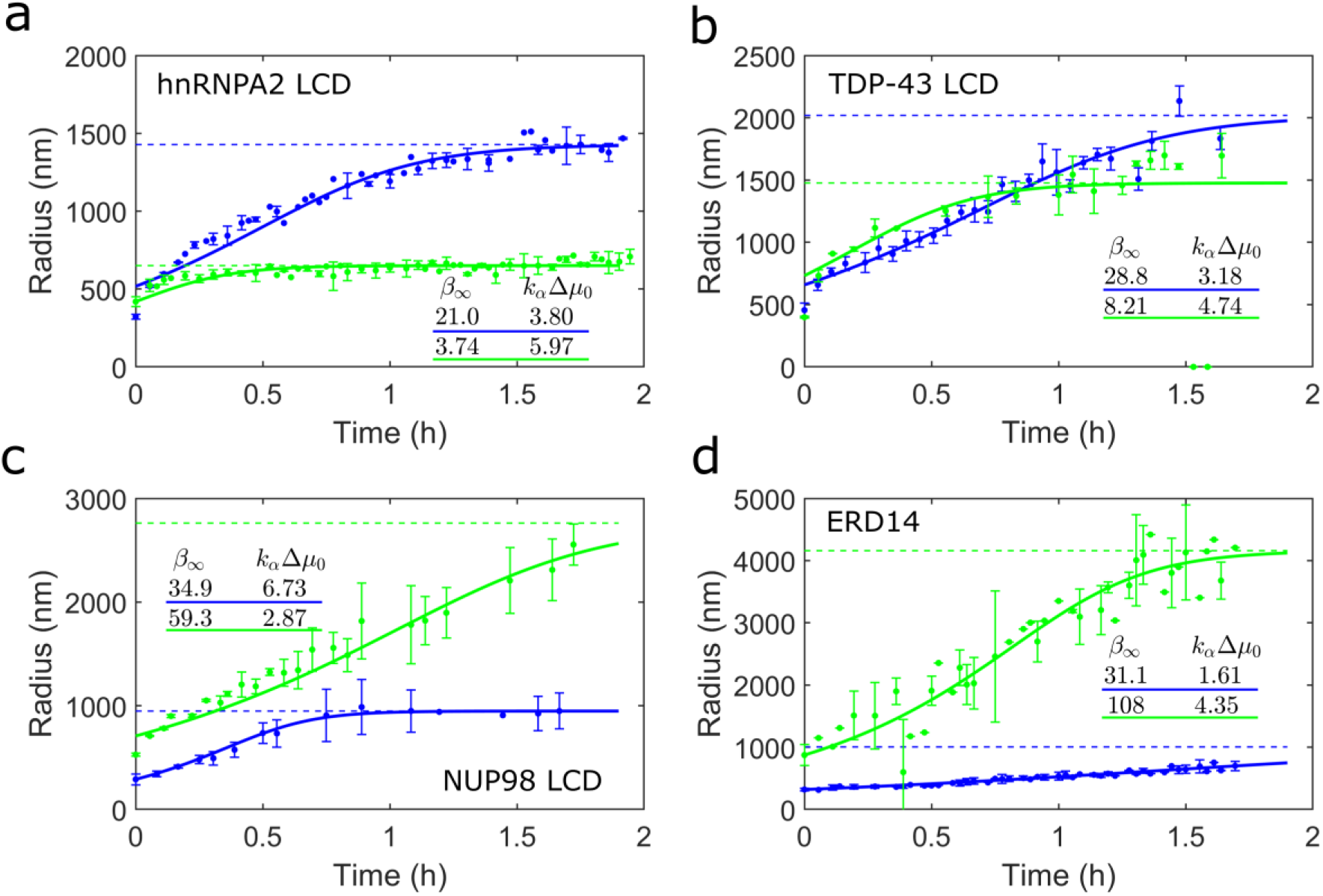
Kinetic analysis of LLPS of proteins (a) 20 μM hnRNPA2 LCD, (b) 80 μM TDP-43 LCD, (c) 10 μM NUP98 LCD and (d) 20 μM ERD14. Symbols: Droplet size evolution measured by Van Lindt et al. [16] in the absence (blue) and presence (green) of 150 mM NaCl [16]. Solid lines: fitted 〈*R*〉(*t*) curve computed as the product 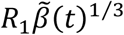 using the values of *β*_∞_ and *k_α_*Δ*μ*_0_ listed in each panel’s table. Dashed lines: predicted value of the final mean size 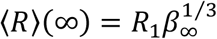.

To ascertain the reasons why larger or smaller droplets are produced in the different experiments it is necessary to know the specific values *α_c_* and *k_β_* that lead to large or small values of *β*_∞_, respectively. In the present example, approximate values of *α_c_* can be determined using estimates of the saturation concentration *c*_∞_ and critical solubility *c_c_* extracted from the 600 nm-turbidity increase that was induced by different protein concentrations [16]. Since LLPS does not spontaneously occur for *c*_0_ < *c_c_*, decreasing values of protein concentration eventually reach a point of *c*_0_ ≈ *c_c_* below which the initial turbidity signal changes no more. Accordingly, values of *c_c_* could be identified (Table 1) from the protein titration results in Figure S9 of the source manuscript [16]. The same experimental data is used to estimate *c*_∞_ values from the x-intercept of the maximum turbidity Δ*F*_max_ vs. *c*_0_ straight line given that the final amount of the dense phase is proportional to the concentration difference Δ*c*_0_ = *c*_0_ – *c*_∞_ [8,9]. As expected for aggregation-prone proteins, very low values of the saturation concentration are obtained for all proteins under study (*c*_∞_ ≈ 0). Once *c*_∞_ and *c_c_* estimates are available, STEs can be quantified as STEs = 1 – *α_c_* using the definition of *α_c_* (Eq. 4). The analysis of the 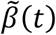 solution in Figs. 3d and S5 shows that 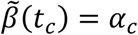 is a good approximation for all combinations of *α_c_* and *k_β_*. Thus, our next step in the characterization of model parameters is the identification of the instant *t_c_* at which the equivalence 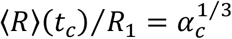 is verified (Fig. 4). Then, the definitions of time constants *τ* and *t*_1_ in Fig. 2a can be expressed as a function of parameters *k_β_*, *α_c_* and *k_α_*Δ*μ*_0_, and replaced in the definition of *t_c_* to obtain the *k_β_* estimate (Table 1). With the *k_β_* results in hand, values of *τ* and *t*_1_ are finally obtained using their mathematical definitions in Fig. 2b.

**Table 1.** Model parameters estimated from the LLPS data measured by Van Lindt et al. [16] and from the fitted *k_α_*Δ*μ*_0_ and *β*_∞_ values in Fig. 4 for the cases of no added NaCl.

| Protein | $c_0$<br>($\mu\text{M}$) | $c_c$<br>($\mu\text{M}$) | $\alpha_c$ (–) | $t_c$ (h) | $R_1$ (nm) | $k_\alpha \Delta\mu_0$ ($\text{h}^{-1}$) | $\beta_\infty$ (–) | $k_\beta$ (–) | $\frac{k_\alpha \Delta\mu_0 \tau}{2}$ (–) |
| --- | --- | --- | --- | --- | --- | --- | --- | --- | --- |
| hnRNPA2<br>LCD | 20 | 2 | 0.90 | 1.333 | 517.2 | 3.80 | 21.0 | 0.074 | 1.01 |
| TDP-43<br>LCD | 80 | 20 | 0.75 | 1.317 | 658.5 | 3.18 | 28.8 | 0.090 | 1.04 |
| NUP98<br>LCD | 10 | 3.8 | 0.62 | 0.598 | 290.2 | 6.73 | 34.9 | 0.086 | 1.04 |
| ERD14 | 20 | 10 | 0.50 | 1.283 | 319.3 | 1.61 | 31.1 | 0.908 | 3.29 |

Of all proteins and conditions considered in Table 1, hnRNPA2 LCD is the one less affected by STEs because the adopted initial concentration is much higher than *c_c_*. In the absence of NaCl, the mean size increase 〈*R*〉(∞)/*R*_1_ experimentally measured for this protein was higher than for ERD14, the protein more strongly affected by STEs - compare Figs. 4a and 4d. This observation is explained by lower nucleation-to-growth ratios (lower *k_β_* values) exhibited by the first protein. However, if LLPS had been followed over a longer period, a ~31-fold increase in droplet volume would be expected for ERD14, a value that is larger than the value of *β*_∞_ obtained for hnRNPA2 LCD (Table 1). The case of ERD14 illustrates the possibility of fast nucleating droplets (*k_β_* = 0.91) that slowly grow into micrometer-sized particles due to the existence of a growth-only period after *t* = *t_c_*. In contrast to what happens for the other proteins, the *τ* values listed in Table 1 for ERD14 are much larger than 2(*k_α_*Δ*μ*_0_)^−1^ as a consequence of more pronounced STEs and of faster nucleation rates. The reason why this protein takes more time to complete LLPS is explained by a slow growth step and by the increased importance of this step in the presence of STEs. As illustrated below in the Discussion section, strong STEs (low *α_c_*) and fast nucleation rates (high *k_β_*) uniquely combine to allow for functional phase separation via sharply regulated responses to minute concentration changes. Interestingly, the combination of stronger STEs (lower *α_c_*) and higher *k_β_* is observed for ERD14 (Table 1), which is an intrinsically disordered chaperone with the function of helping plants to survive under dehydration stress [39,40], whereas hnRNPA2, TDP-43 and NUP98 occur in bimolecular condensates associated with disease [16,41].

### Predictive Power

We now investigate whether the two-parameter physical model can quantitatively predict the evolution of size distributions during the phase separation of TDP-43 LCD and NUP98 LCD. For these proteins, the measured DLS autocorrelation functions (Fig. S6) yield reliable size distributions in all stages of LLPS (Figs. 5a and 5b). To compare the experimental data with the model predictions, light scattering intensities are simulated as a function of time and droplet size using the zeroth *P*(*t*), first *M*(*t*) and second *Q*(*t*) moments of the size distribution, from which the mean size *N*(*t*) (first cumulant) and the variance *σ*^2^(*t*) (second cumulant) are determined (Appendix: *Size Distributions*). This strategy is an alternative to the recursive integration of the master equation (Eq. S2) over the possible values of *j* = *N*_1_, *N*_1_ + 1,…, *N*_∞_, a method that becomes inpractical for the droplet sizes in question.

**Figure 5.**
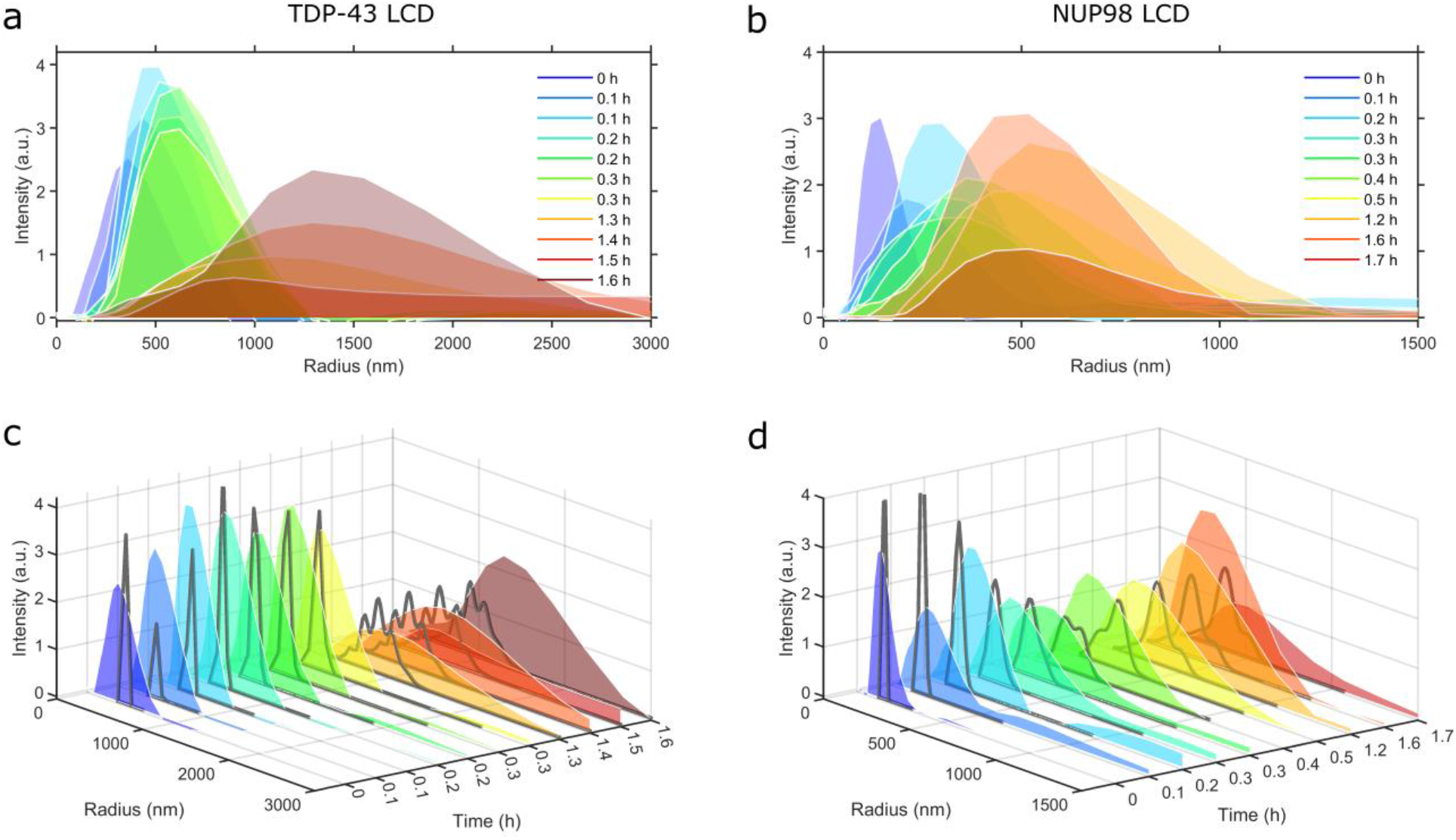
Time variation of the droplet size distributions. (a and b) DLS intensity distributions obtained for (a) TDP-43 LCD and (b) NUP98 LCD. (c and d) Gray lines: light scattering intensities predicted by the general nucleation-and-growth model for (c) TDP-43 LCD and (d) NUP98 LCD using the values of *α_c_*, *k_α_*Δ*μ*_0_ and *k_β_* listed in Table 1. Further numerical details in *SI Additional Methods*. Different colored curves: same experimental results as in (a) and (b).

The computed curves follow a similar trend to the one observed for the mean size, shape and scattering intensity of the DLS distributions (Figs. 5c and 5d). This fact is all the more remarkable when we consider that no numerical adjustments were required besides the choice of a fixed proportionality factor normalizing the scattering intensities during each experiment and that automatic Mie scattering corrections are applied to the simulated size distributions. Moreover, the droplet sizes are markedly distinct in the cases of TDP-43 LCD and NUP98 LCD and spread over distinct range values as LLPS proceeds. The combination of results in Figs. 4 and 5 illustrates that the general nucleation-and-growth model has the potential to characterize, and even predict intricate aspects of phase separation using the *α_c_* measurable and two kinetic parameters.

## Discussion

The sub-critical region of the phase diagram opens surprising avenues for deciphering the origins of intriguing biological processes, from pathogenic protein aggregation to the spatiotemporal control of condensate formation in the cell. Proteins can be stable and functional under supersaturated conditions without the immanent risk of undergoing primary nucleation and forming potentially toxic aggregates. In this light, the association of protein misfolding disorders with ageing would not result from the fact that critical macromolecules can be naturally supersaturated [42], but would rather reflect the increasing chances of the critical supersaturation barrier being crossed as the individual grows old. Such transgressions are likely to occur as the result of, for example, incidental cell stress, chemical modifications of the protein and altered proteostasis. If the nuclei resist cellular quality control mechanisms, a single species might be enough to elicit a cascade of events comprising the secondary nucleation of new aggregates and the cell-to-cell spreading of the disease in a prion-like fashion [43]. Another possibility opened by STEs is the formation of large and uniformly distributed particles without recourse to coalescence or external players such as enzymes and chaperones. This possibility was hard to conceive from a purely biophysical point of view: either primary nucleation is fast and steady-state conditions are rapidly achieved through the formation of numerous small particles, or primary nucleation is slow relative to the growth step and non-uniform size distributions are obtained with very large particles coexisting with very small ones (Figs. 1c and 1d, Movie S1). The occurrence of large and uniformly distributed condensates in the cell has been enigmatic and is often ascribed to some unknown or not fully outlined regulatory process(es). However, the observation of the same pattern in test-tube experiments demonstrates that STEs are essential for the understanding of biological phase separations. Narrow size distributions are expected in the presence of strong STEs because the occurrence of primary nucleation becomes limited to the initial moments of phase separation (Figs. S3c and S3d and Movie S2). Splitting nucleation-and-growth into two consecutive steps facilitates the production of large condensates with precisely controlled size distributions.

By deliberately simplifying our analysis to phase separation of binary mixtures, we demonstrate that size heterogeneity is not the inexorable result of nucleation-and-growth processes which, on the contrary, can be spontaneously regulated according to basic physicochemical principles of phase equilibrium. This said, considering STEs at the quantitative level is particularly useful for understand how multicomponent LLPS can be accurately controlled in the cell through the modulation of interfacial properties. Being interfacially active, crucial proteins, such as G3BP1 in stress granules, MEG-3 in P granules, centrosomin and DSpd-2 in the centrosome, and TPX-2 in the mitotic spindle, can induce a generalized solubility drop in the lighter phase that provokes the recruitment of all molecule whose concentration is above the critical limit. Surface tension modulation, therefore, emerges as a general organizing principle regulating phase separations in health and disease.

We envisage a practical application of our biophysical model in drug discovery, namely on the quantification of the effects of STEs-modulators and kinetic inhibitors on the progress curves of biological self-assembly. These curves can be measured in the presence and absence of drug candidates and then fitted to the exact *α*(*t*) solution (Fig. 2a) or the approximate 〈*R*〉(*t*) solution (Fig. 3a) depending if mass-based or size-based data are available. Deviations from the theoretical 〈*R*〉(*t*) curve (Fig. 3a) are expected if significant secondary nucleation occurs. In that case, the exact numerical solution of Eq. 5 or the approximate 〈*R*〉(*t*) solution assuming no STEs present [9] can be adopted in the alternative. In principle, the *α*(*t*) solution (Fig. 2a) is widely applicable independently of the magnitudes of STEs and secondary nucleation rates. Size distribution analysis may additionally reveal the continuum transition from the monomeric state, to mesoscale clusters, to the formation of a new liquid phase [44]; when this happens, sharp nucleation-and-growth models do not apply and should be replaced by oligomerization equilibrium analysis.

Quantifying the effect of kinetic inhibitors on the rates of growth, primary nucleation, and secondary nucleation is well-established in protein misfolding research [7,45], but it is almost absent from biological LLPS research where chemical kinetic analysis is generally based on semi-empirical scaling laws and exponents. In addition to the *α*(*t*) and 〈*R*〉(*t*) solutions here derived, we also proposed novel methodologies to quantify STEs from the measurements of the saturation concentration *c*_∞_ and critical solubility *c_c_* (see analysis to Fig 4). We showed, moreover, that the discrete size distributions of biological condensates can be inferred at the quantitative level using kinetic parameters fitted to *α*(*t*) or 〈*R*〉(*t*) progress curves (Fig. 5). This opens new possibilities for gauging the effect of phase separation modulators from measured values of equilibrium concentration and size-distribution parameters.

## Conclusions

Biological phase separation is associated with multiple aspects of cell physiology and may be conducive to neurodegenerative processes. How, when and where this phenomenon occurs remains difficult to predict even under test-tube experimental conditions. Here we propose a physical description of protein phase separation that explains intriguing observations such as the spontaneous formation of a few condensates with large and uniform sizes. We show that the caveats in the current nucleation-and-growth theory can be solved by addressing the role of surface tension during the formation of primary nuclei. Shape- and size-dependent STEs provoke an increase in the thermodynamic solubility, which is reflected in higher energetic barriers for primary nucleation. A “margin of safety” is therefore introduced so that the birth of the new phase becomes thermodynamically unfavored in metastable regions of the phase diagram. The difference between the saturation limit and the critical solubility can be interpreted in the light of the Young-Laplace equation for curved interface equilibria. These principles apply to liquid-solid and LLPS alike and are included in the master equation describing nucleation-and-growth phenomena (Eqs. 5 and S2), whose closed-form solution is here derived (Fig. 2a). In most situations, the exact solution simplifies to an exponential-type function describing how the average particle size changes over time (Fig. 3a). The model predictions are validated using kinetic data measured for different proteins during the phase separation of different-sized biological condensates. We found that the final size, as well as the moment of formation of biological condensates become highly regulated in the presence of strong STEs and that the evolution of droplet size distributions can be predicted based on the *α_c_* measurable and two kinetic parameters (Figs. 5c and 5d). The generalized nucleation-and-growth model can be used for chemical kinetics analysis of isolated systems, but it could also be extended over complex, multicomponent systems for interrogating mechanisms of cell regulation and neurotoxicity involving phase separation.

## Materials and Methods

### Model Validation

The general nucleation-and-growth model and its validity are tested using kinetic data of LLPS (Fig. 4) and amyloid aggregation (Fig. S1), by rationalizing the formation of phase-separated condensates with large and uniform sizes under conditions where active biological processes are not present (Fig. S3d and Movies S1 and S2), and by comparing the predicted time evolution of droplet sizes with the measured one (Figs. 5c and 5d). The raw data for graphs presented in Fig. 4 are available in the Supplementary Data files of reference [16]. The raw data presented in Fig. S1 were digitized by us from the original reference [10]. Details of the adopted numerical methods in *SI Additional Methods*.

### DLS measurements

Measurements of DLS autocorrelation functions were carried out on a DynaPro NanoStar (Wyatt) instrument during LLPS of 80 μM TDP-43 LCD and 10 μM NUP98 LCD induced by pH jumps from 3.5 to 7.5 (TDP-43 LCD) and 3.0 to 7.5 (NUP98 LCD). A disposable cuvette (WYATT technology) was filled with 100 μl of protein solution at pH and concentration values at which LLPS occurred. The sides of the cuvette were filled with water and a cap was put on top. The samples were illuminated by a 120 mW air-launched laser of 658 nm wavelength intensity and the scattered light was recorded at a scattering angle of 90° at 25 °C, for a period of 6 h, collecting 10 acquisitions (8 s each). Each measurement was repeated at least 3 times. Hydrodynamic radii of the particles in solution were estimated from the diffusion coefficient delivered from the analysis of measured autocorrelation functions as described in Figs. S6a and S6b.

## Supporting information

Simulated time-lapse of particle self-assembly using the same model parameters as described in Fig. 1d.

Simulated time-lapse of particle self-assembly using the same model parameters as described in Fig. S3d.

## Supporting Information

- Appendix

- Additional Methods

- Additional Discussion

- Additional Figures

- SI References.

- Movies S1 and S2

## Acknowledgements

This work is part of a project that has received funding from the European Union’s Horizon 2020 research and innovation programme under grant agreement No. 952334 (PhasAGE). This research was financed by (i) FEDER—Fundo Europeu de Desenvolvimento Regional funds through the COMPETE 2020—Operational Programme for Competitiveness and Internationalization (POCI), Portugal 2020, and by Portuguese funds through FCT—Fundação para a Ciência e a Tecnologia/ Ministério da Ciência, Tecnologia e Ensino Superior (FCT/MCTES) in the framework of projects PTDC/QUI-COL/2444/2021, POCI-01-0145-FEDER-031323 (PTDC/MED-FAR/31323/2017), and POCI-01-0145-FEDER-007274 (“Institute for Research and Innovation in Health Sciences”), (ii) by Base Funding–UIDB/00511/2020 of the Laboratory for Process Engineering, Environment, Biotechnology and Energy (LEPABE) funded by national funds through the FCT/MCTES (PIDDAC).

## Supporting Information (SI)

- Appendix
  ○ General Nucleation-and-Growth Model
  ○ Size Distributions
- Additional Methods
  ○ Numerical Methods
- Additional Discussion
  ○ Kinetic Analysis of Amyloid Aggregation
  ○ STEs on phase diagrams and particle size distributions
- Additional Figures
  ○ Figure S4. The impact of secondary nucleation on the final particle size in the absence of STEs.
  ○ Figure S5. Maximum error of the 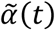 and 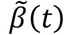 approximations (Fig. 3a) relatively to the exact solution of Eq. 5.
  ○ Figure S6. DLS autocorrelation functions during LLPS of (a) TDP-43 LCD and (b) NUP98 LCD.
  ○ Figure S7. Using the gamma distribution to describe the exact solution of the master model equation (Eq. S2).
- SI References.

## Appendix

### General Nucleation-and-Growth Model

In previous master equations describing nucleation-and-growth of amyloid fibrils, the concentration of filaments composed of *j* ≥ *N*_2_ monomers (*C_j_*) is determined by the rate constants *k*_1_, *k*_2_, *k*_+_ and *k*_−_ characterizing the processes of primary and secondary nucleation, growth, and fragmentation, respectively [1,2]:

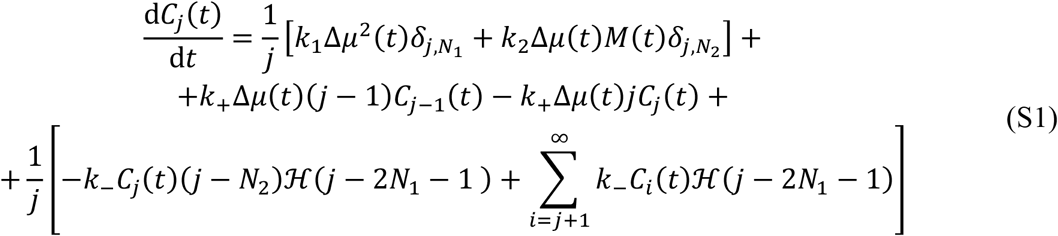

where the Kronecker delta function sets the sizes of the primary and secondary nuclei and the Heaviside function establishes a minimum size of 2*N*_1_ + 1 molecules above which fragmentation starts to occur [2]. We assume that STEs mainly affect the driving force for primary nucleation and thus only the *k*_1_ term has to be expressed in relation to the critical supersaturation level (Δ*μ_c_*):

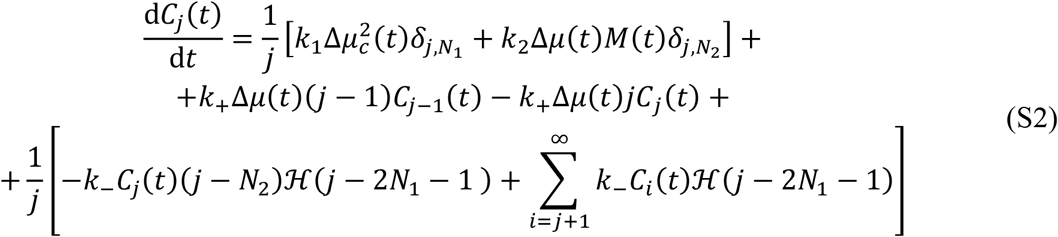

The number and mass concentrations correspond to the zeroth and first moments of the *C_j_*(*t*) distribution:

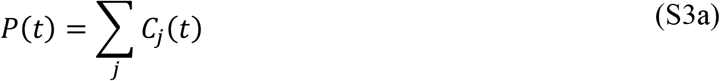

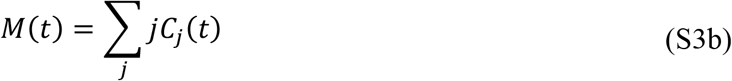

and are used to obtain Eqs. 5a and 5b after differentiation, replacement of Eq. S2 and algebraic manipulation of the summations; to obtain Eq. 5a, negligible fragmentation rates are assumed, whereas in Eq. 5b the terms dependent on *k*_−_ ultimately cancel each other. Furthermore, from the definitions of *α*(*t*) = *M*(*t*)/*M*_∞_, *M*_∞_ = *c*_0_ – *c*_∞_ and Δ*μ_c_*(*t*)/Δ*μ*_0_ (Eq. 3), constants *k_α_* and *k_β_* can be defined as a function of the elementary rate constants as follows:

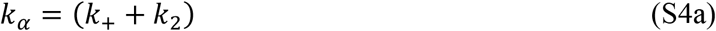

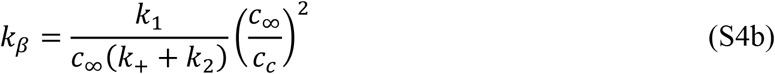

In the absence of STEs, *c_c_* = *c*_∞_ and Δ*μ_c_* = Δ*μ*, which simplifies Eqs. 5a and 5b to a system of ODEs of known analytical solution [1, 2]:

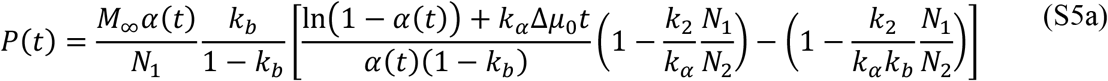

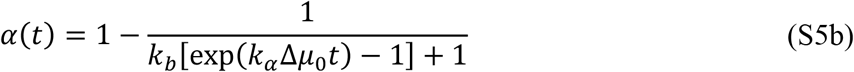

where *k_b_* = *k_n_*/(*k*_+_ + *k*_2_), with *k_n_* = *k*_1_/*c*_∞_ [1, 2].

In the presence of STEs, Eq. 5b can be solved isolatedly to obtain the following analytical solution:

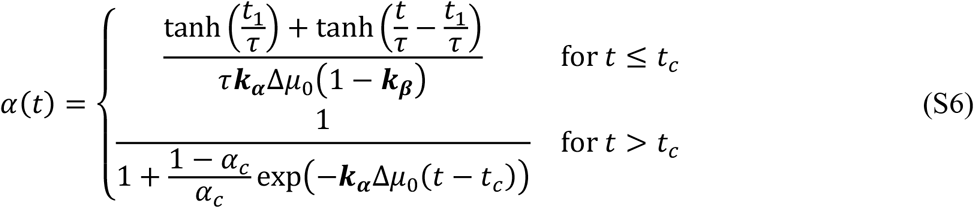

where

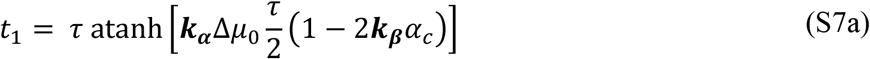

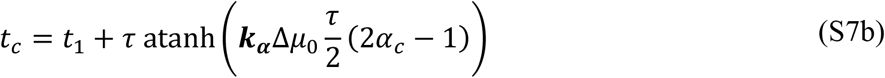

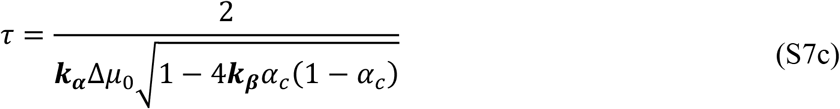

Therefore, Eq. S5b is the limit case of Eq. S6 when *α_c_* = 1 (no STEs present).

### Size Distributions

The particle size distributions (PSDs) predicted by the general nucleation-and-growth model are obtained by recursive integration of Eq. S2 over the possible values of *j*. This method cannot be always adopted since heavy computational requirements are required when larger particles are considered: in the exemplary case of a 10 kDa globular protein, a small spherical droplet of ~100 nm radius would contain *j* > 3 × 10^5^ monomeric units. As an alternative, the mean size *N*(*t*) and variance *σ*^2^(*t*) of the *C_j_*(*t*) distribution can be determined as a function of the principal moments using the definitions of the first and second cumulants:

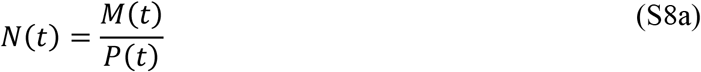

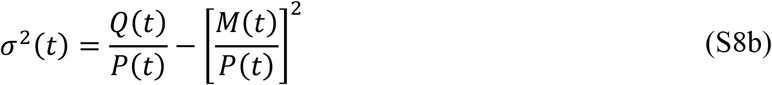

The definition of the second moment,

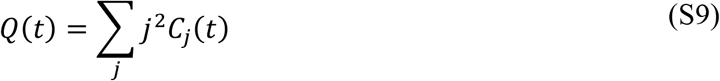

is differentiated and used together with Eq. S2 and the *P*(*t*), *M*(*t*) sumations (Eq. S3) to obtain:

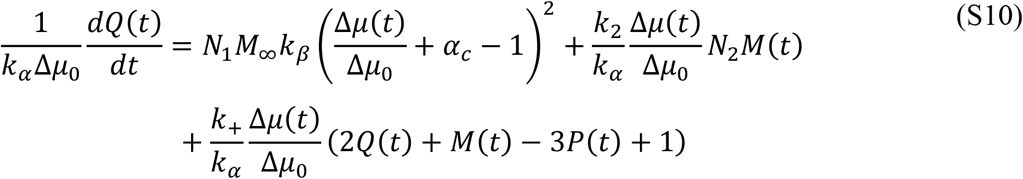

Therefore, an exact solution for the zeroth, first and second moments is possible by solving Eqs. 5a, 5b and S10 simultaneously. As described in detail in *SI Additional Methods*, the mean and variance of *C_j_*(*t*) are used to predict the shape of droplet size distributions measured by DLS (Figs. 6c and 6d of the main text).

## Additional Methods

### Numerical Methods

The model equations are fitted to experimental data in Figs. 4 and S1 using the *lsqcurvefit* function of Mathworks MATLAB R2020b (Natick, MA, USA). The PSDs in Figs. 1d and S3d are generated by numerically solving Eq. S2 for long reaction times using the model parameters identified in each figure’s caption assuming that *N*_1_ = 2 monomers, that no condensates are initially present *C_j_*(0) = 0, and a total of *M*_∞_ = 1000 monomers separating into the new phase. Each particle size distribution is normalized by the maximum frequency value and represented as a function of *R_j_*/*R*_1_ = (*N_j_*/*N*_1_)^1/3^. In Movies S1 and S2, the number of condensates represented in each timeframe coincides with the total number of particles calculated per unit volume; condensates composed of *N_j_* monomers are schematically represented as circles with symbol size *N_j_*.

To compute the *α*(*t*) progress curves (Figs. 1c and S3c), the system of ordinary differential equations (ODEs) comprising Eqs. 5 and Eq. S2 was numerically solved and the results expressed in *k_α_*Δ*c*_0_-normalized units of time.

The theoretical size distributions in Figs. 6c and 6d are generated from the solution of Eqs. 5a, 5b and S10 for the values of the *α_c_*, *k_α_*Δ*μ*_0_ and *k_β_* parameters listed in Table 1 for TDP-43 LCD and NUP98 LCD. Negligible secondary nucleation (*k*_2_ ≈ 0) and the absence of initial condensates are assumed; the used *R*_1_ values are inferred from the initial DLS distribution (450 nm for TDP-43 LCD and 160 nm for NUP98 LCD). Replacing the *P*(*t*), *M*(*t*) and *Q*(*t*) solutions in Eq. S8 gives the variation with time of the mean size and variance of the distributions. We verified that the gamma probability distribution adequately describes the gradual change from exponential to normal distribution predicted by the master model equation Eq. S2 (Fig. S7). The approximate *C_j_*(*t*) profiles are obtained from the gamma distribution function using as parameters *θ*(*t*) = *σ*^2^(*t*)/*N*(*t*) and *k*(*t*) = *N*(*t*)/*θ*(*t*) [3]. Finally, each droplet is modelled as a Mie scatterer, whose scattering intensity is calculated from the complex scattering amplitudes for two orthogonal directions of incident polarization. The MATLAB code *MatScat* [4] was used for this purpose assuming: a spherical geometry, collection of scattered light between scattering angles of 80 and 100 degrees, a complex refractive index of 1.39 + 0.01*i*, a outer-medium refractive index of 1.333, a wavelength of 658 nm, and a proportionality factor of 1.11 × 10^16^ (TDP-43 LCD) and 1.00 × 10^16^ (NUP98 LCD) to normalize scattering intensity units.

## Additional Discussion

### Kinetic Analysis of Amyloid Aggregation

The kinetics of amyloid aggregation of monomeric transthyretin (mTTR) was fully characterized by Hurshman et al. from the increase of 400 nm-turbidity and thioflavin-T fluorescence over time [5]. Only the measurements using the amyloid-specific dye are here used to quantitatively analyze mass-based *α*(*t*) progress curves (Fig. S1a). In previous analyses of TTR aggregation kinetics, the experimental data was well fitted by values of *k_β_* ≫ 0.01, which, nevertheless, changed with protein concentration in a manner not expected by the Classical Nucleation Theory (CNT) [1]. The applied concentrations of mTTR between 50 and 400 μg/mL are much higher than the solubility value of *c*_∞_ = 0.8 μg/mL that was estimated from the monomer concentration in solution after long reaction times (1 week) independently of the departing mTTR concentration [5]. Although the terms *protein solubility* and *protein critical concentration* are often adopted interchangeably in the literature, the occurrence of STEs demands a clear distinction between *c*_∞_ and *c_c_*. The data measured by Hurshman et al. does not provide a direct estimate of *c_c_* because the thioflavin-T signal becomes insensitive to protein aggregation for mTTR concentrations below 5 μg/mL [5]. Values of *c_c_* between *c*_∞_ = 0.8 μg/mL and 5 μg /mL are therefore acceptable. Model parameters *k_α_* and *k_β_*, and the additional unknown *c_c_* can be numerically fitted without overparameterization problems from the analysis of the effect of protein concentration on phase separation kinetics (Fig. S1a). The obtained value of *c_c_* = 4.06 μg/mL implies that the magnitude of STEs is low: *α_c_* = 0.93 from Eq. 4 using as *c*_0_ the lowest concentration tested (50 μg/mL). Although only mild STEs are present, their contribution allows to adopt a single pair of concentrationin-dependent kinetic constants to globally fit the measured effects of mTTR concentration on the *α*(*t*) time-course curves (Fig. S1a).

**Figure S1.**
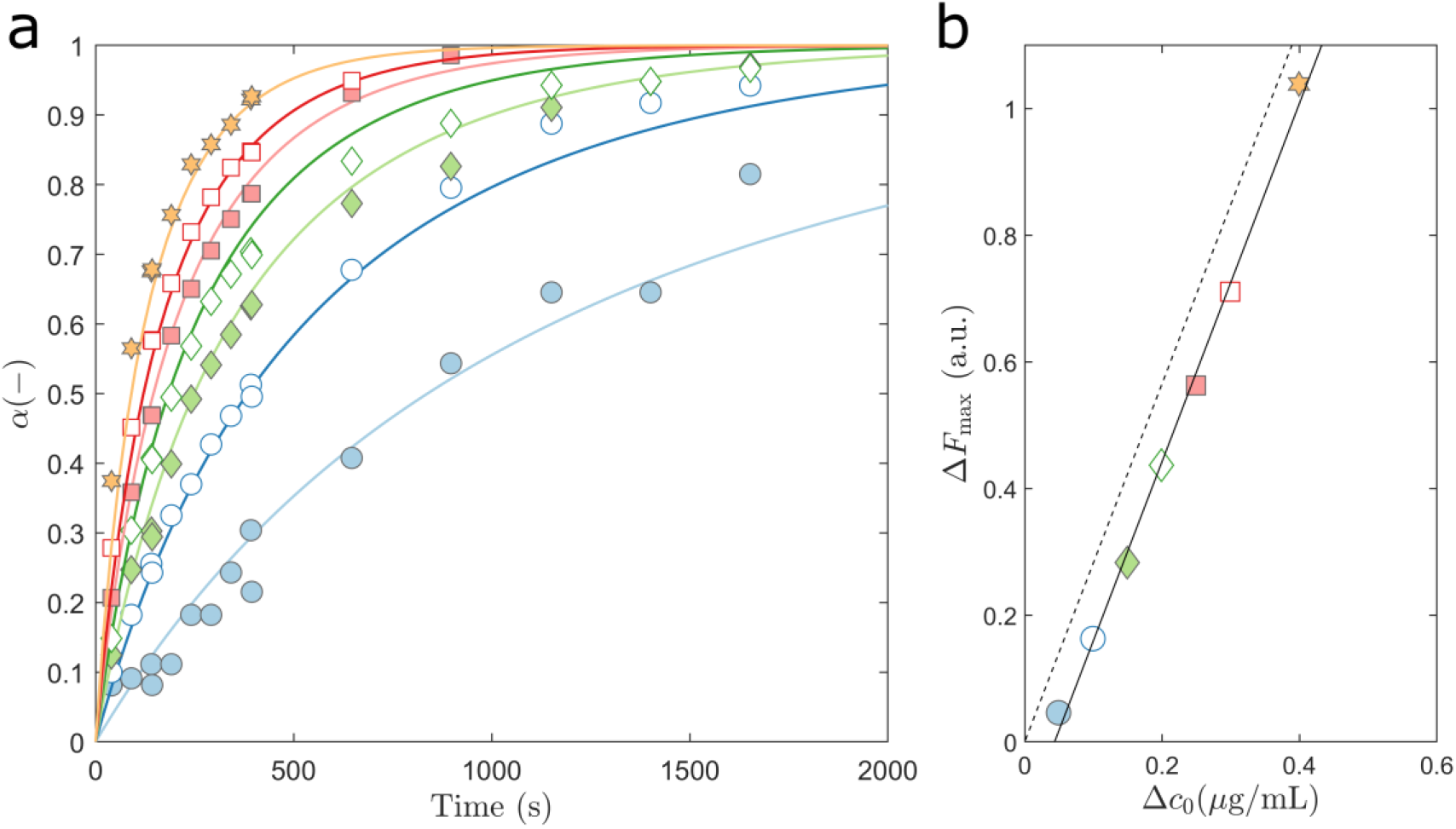
The general nucleation-and-growth model describes the (a) kinetics and (b) thermodynamics of mTTR aggregation measured by Hurshman et al. [5]. (a) Symbols: continuous fluorescence data digitized by us into periodic data and normalized by the end-point signal for mTTR concentrations of (from top to bottom) 400, 300, 250, 200, 150, 100 and 50 μg/mL. Solid lines: Progress curves predicted by Eq. S6 for the tested mTTR concentrations using the fitted parameters *k_α_* = 1.05 × 10^−2^ s^−1^, *k_β_* = 1.71 and *c_c_* = 4.06 μg/mL. (b) Symbols: Values of maximum fluorescence measured as a function of Δ*c*_0_ = (*c*_0_ – *c*_∞_). Solid line: the experimental results follow a linear correlation. Dashed line: example of a straight line passing in the origin.

To cross-check the validity of protein-aggregation kinetic analyses, the maximum fluorescence signal obtained at the end of the reaction Δ*F*_max_ should be evaluated against the concentration difference Δ*c*_0_ = (*c*_0_ – *c*_∞_). A direct proportion between these two quantities is expected in the absence of parallel aggregation pathways [1,6]. While a linear relationship between Δ*F*_max_ and Δ*c*_0_ is confirmed, the positive X-intercept suggests the occurrence of TTR in other forms than monomers or amyloid fibrils (Fig. S1b). Confirming this indication, significant amounts of TTR dimers were identified by Hurshman et al. using analytical gel filtration and SDS-PAGE techniques [5]. Overall, the results in Fig. S1 confirm that the effect of protein concentration on mTTR aggregation is well described by standard nucleation theories modified to include STEs.

A distinguishing feature of the general nucleation-and-growth model is the predicted effect of protein concentration on the shape of reaction progress curves. Expectedly, in the case of fast-nucleating proteins such as mTTR, lowering the initial concentration not only slows down the rates of PS as it will also change the shape of the *α*(*t*) curves from hyperbolic to sigmoidal. This new possibility is admissible when the time interval during which primary nucleation occurs is significantly reduced by the occurrence of strong STEs. As we have seen for mTTR aggregation, only weak STEs are present in the range of protein concentrations studied by Hurshman et al. [5]. If the formation of amyloid fibrils of mTTR could be detected accurately for concentrations lower than 50 μg/mL, magnitudes of STEs higher than 0.07 (*α_c_* < 0.93) could be explored, and a lag period of apparently no aggregation would become visible as *c*_0_ gradually approached the value of *c_c_ = 4.1* μg/mL (Fig. S2, dashed lines). Based on this property, it is possible to gauge the intensity of STEs by graphically representing the reaction conversion as a function of Δ*c*_0_-normalized units of time: in principle, superimposing *α*(Δ*c*_0_*t*) curves indicate that STEs are either weak or absent (Fig. S2).

**Figure S2.**
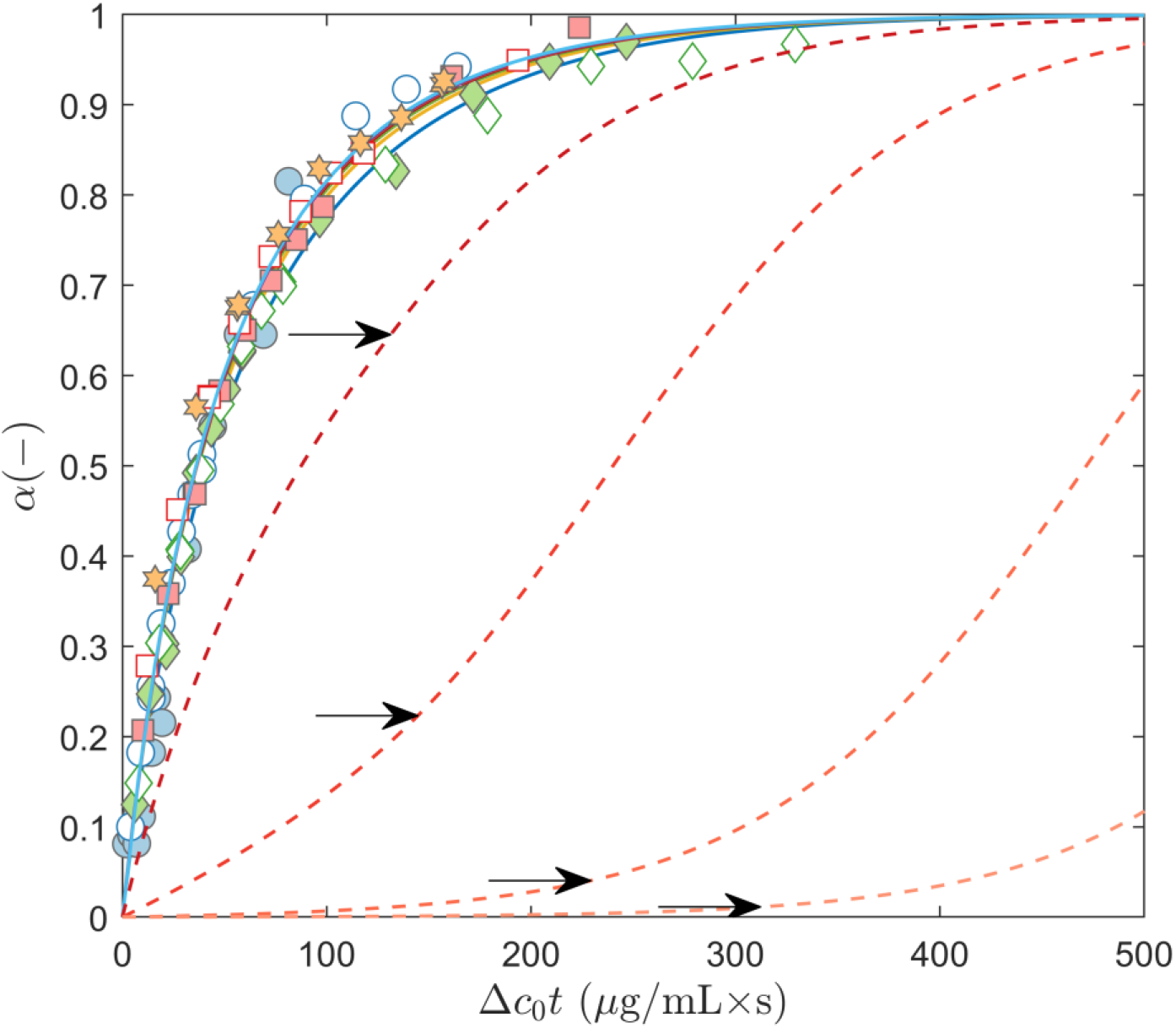
Identification of STEs from the representation of progress curves in Δ*c*_0_-normalized units of time. Symbols and solid lines: the results of Fig. S1a are now represented in normalized time units. Weak STEs influencing mTTR aggregation are confirmed by superimposing *α*(Δ*c*_0_*t*) curves. Dashed lines: predicted *α*(Δ*c*_0_*t*) curves for *c*_0_ values of (from left to right) 10, 5, 4.2 and 4.1 μg/mL using the fitted parameters *k_α_* = 1.05 × 10^−2^ s^−1^, *k_β_* = 1.71 and *c_c_* = 4.06 μg/mL. Arrows: critical reaction conversions (*α_c_*).

### STEs on phase diagrams and particle size distributions

By including STEs in the theory of primary nucleation, a thermodynamic explanation emerges for the recurrent observation of a critical supersaturation limit below which phase separation does not occur at all unless seeding material is added. In terms of the CNT, this limit has been interpreted in relation to awaiting times arbitrarily chosen, below which nucleation is practically arrested [7]. In the presence of STEs, a thermodynamic (rather than kinetic) metastable zone exists where infinitely long times are required for primary nucleation to take place (in blue in Figs. S3a and S3b). In this sense, the width of the metastable zone is a parameter that can be objectively measured and is independent of the method applied to follow phase separation. Without the “margin of safety” created by STEs, protein self-assembly would be almost inevitable in the metastable environment of the cell [8].

**Figure S3.**
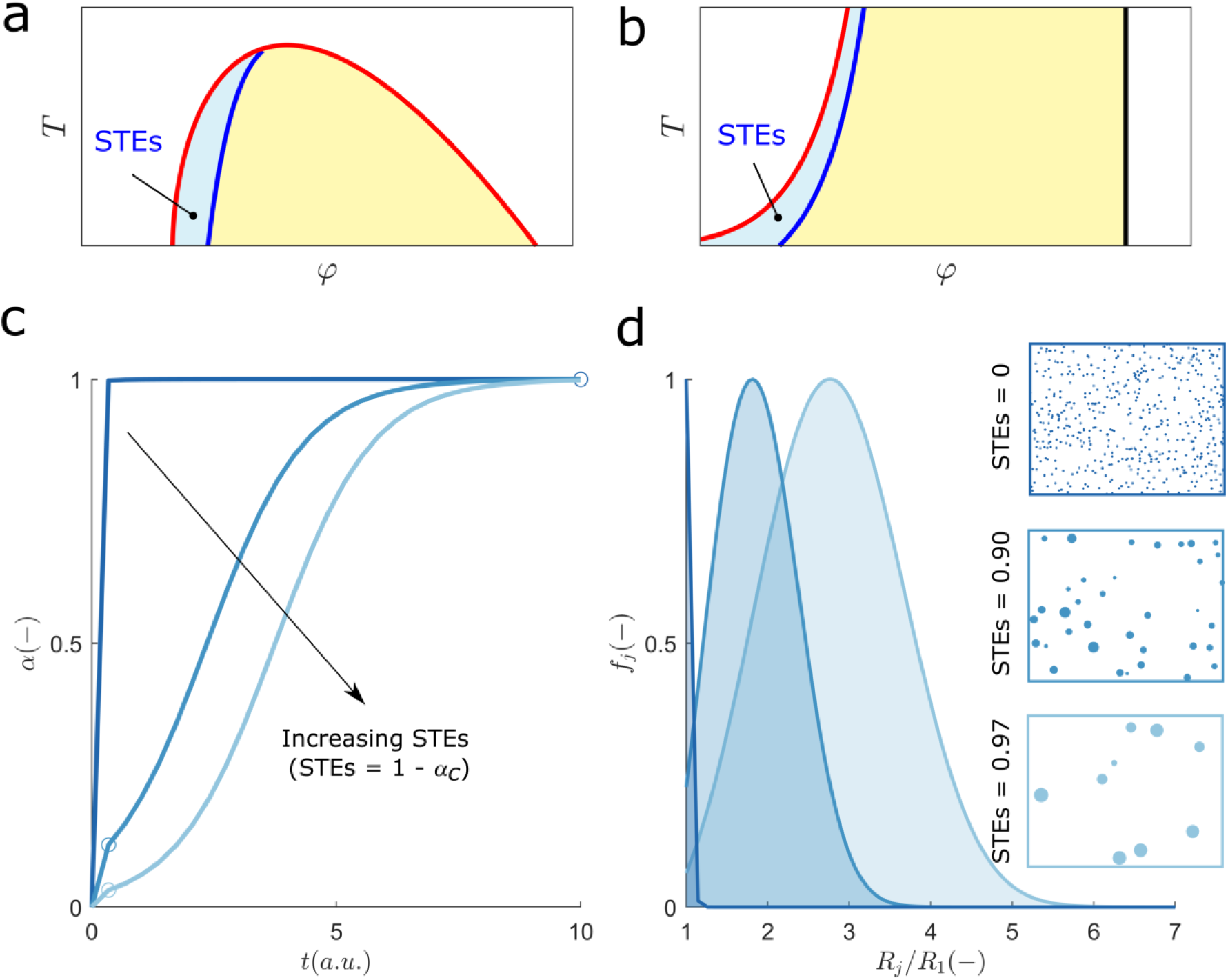
Thermodynamics and kinetics of nucleation-and-growth affected by STEs (refer to Fig. 1 of the main text for the case of no STEs present). (a and b) In the presence of STEs, primary nucleation becomes thermodynamically unfavored in the metastable region (blue area) of (a) liquid-liquid and (b) liquid-solid phase diagrams. (c) Theoretical progress curves calculated using *k_β_* = 10^3^ and (from left to right) STEs = 0, 0.90 and 0.97. Open circles: location of the critical coordinates *t_c_* and *α_c_*. (d) Final *f_j_* distribution calculated using the same model parameters and colour code as in (c). Large particle sizes uniformly distributed around the mean size can be produced in the presence of strong STEs (compare with Fig. 1d). Further numerical details in *SI Additional Methods*.

In the presence of strong STEs, a long growth-only period follows the short nucleation-and-growth period (Fig. S3c) giving rise to a small number of large particles (Fig. S3d). As we tend to the extreme of STEs = 1, the initial formation of a single nucleus would be sufficient to lower the supersaturation level below the critical limit and phase transition would produce only one very large condensate. Movies S1 and S2 are time-lapse simulations showing how particles self-assemble until reaching the equilibrium size distributions in Fig. 1d and S3d, respectively. From comparing the two situations, we conclude that the combination of fast nucleation with strong STEs (movie S2, STEs = 0.97) gives rise to a few large particles and, at the same time, leads to a sharp control of the moment new particles are formed. These features of spatiotemporal regulation of phase separation are not possible in the case of no STEs present: although large condensate bodies can still be produced under nucleation-limited conditions (Fig. 1d), new nuclei will keep arising erratically throughout the whole duration of phase separation (movie S1, *k_β_* = 5 × 10^−3^).

## Additional Figures

**Figure S4.**
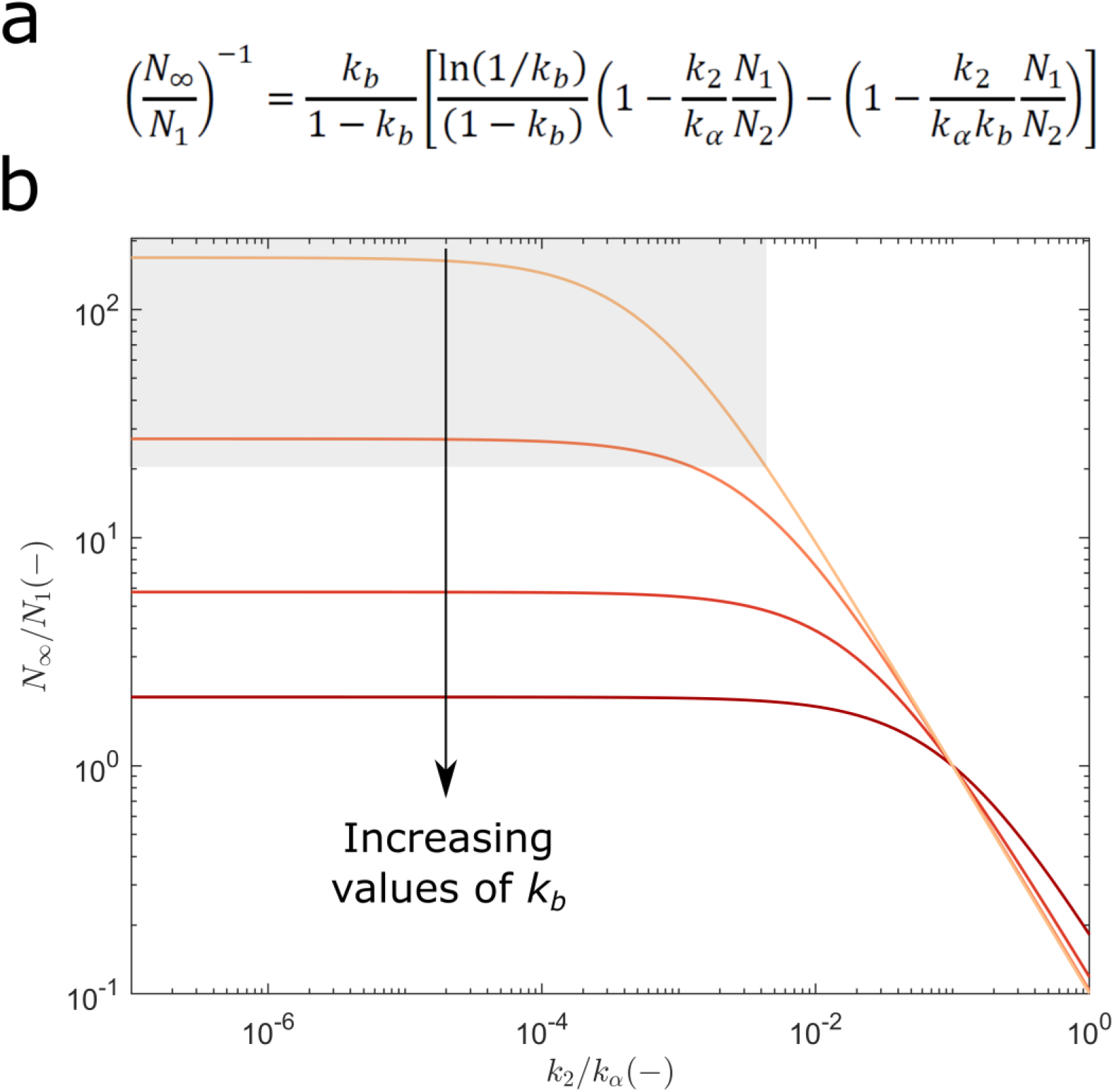
The impact of secondary nucleation on the final particle size in the absence of STEs. (a) The steady-state average particle size *N*_∞_ can be calculated from Eqs. 5c and 5b extrapolated for long reaction times (*t* → ∞). (b) The *N*_∞_/*N*_1_ ratio is calculated as a function of the *k_α_*-normalized value of *k*_2_ for values of *k_b_* of (from top to bottom) 10^−3^, 10^−2^, 0.1, and 1 using *N*_2_/*N*_1_ = 0.1. Primary and secondary nucleation have to be much slower than growth for large particles to be formed (i.e., low values of both *k_b_* and *k*_2_/*k_α_* are required to reach the shaded area).

**Figure S5.**
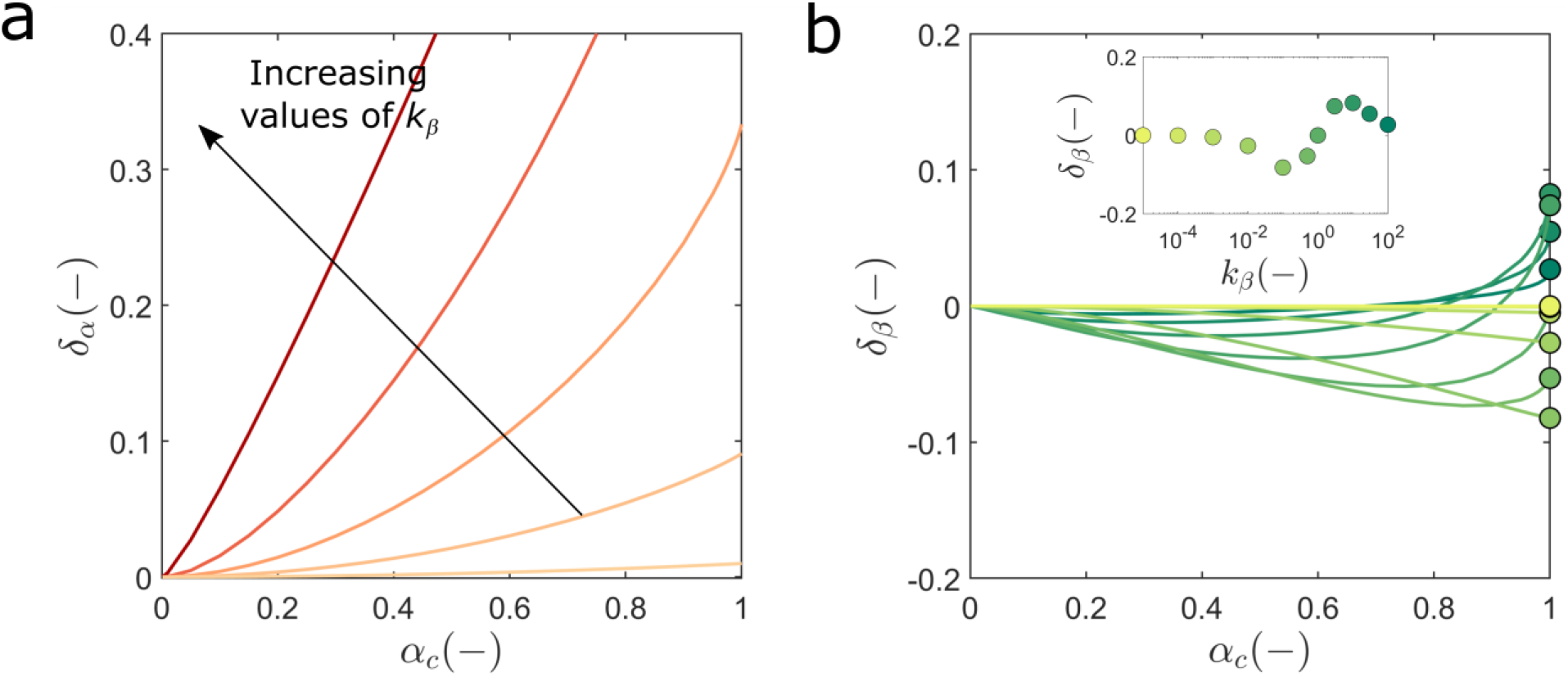
Maximum error of the 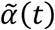 and 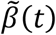 approximations (Fig. 3a of the main text) relatively to the exact solution of Eq. 5. (a) Initial difference 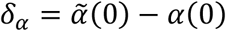 for unseeded reactions (*α*(0) = 0). The estimates of 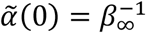 are calculated from the definitions of *β*_∞_ (Fig. 3b) and *t_c_* (Fig. 2b) for values of *k_β_* of (from lighter to darker shades of red) 10^−2^, 10^−1^, 0.5, 3 and 10^2^. (b) Initial difference 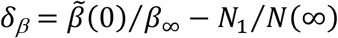, where 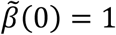 (Fig. 3a) and *N*_1_/*N*(∞) is numerically calculated for values of *k_β_* from (lines from lighter to darker shades of green) 10^−5^ to 10^2^. Symbols: higher values of *δ_β_* are obtained for *α_c_* values close to 1. Inset: the selected *δ_β_* values are represented as a function of *k_β_*.

**Figure S6.**
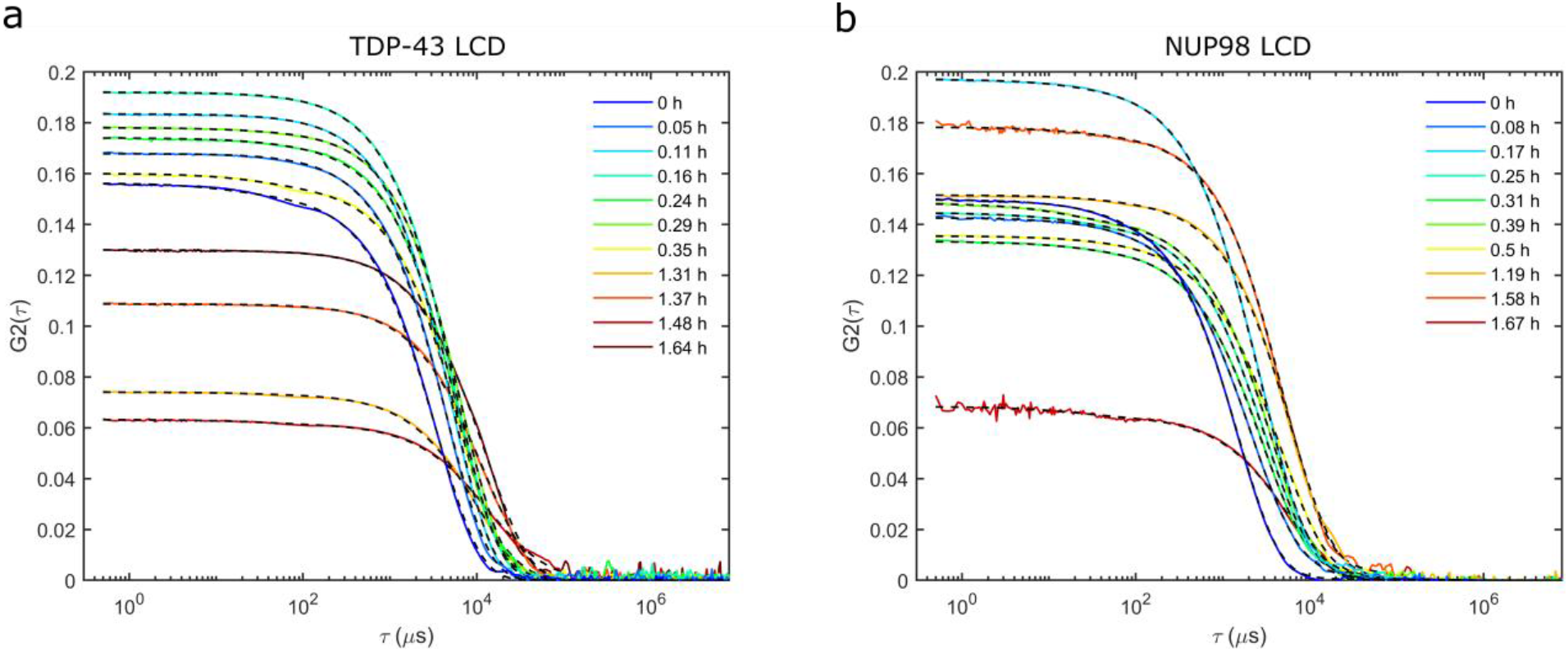
DLS autocorrelation functions during LLPS of (a) TDP-43 LCD and (b) NUP98 LCD. Solid lines: measured data from which the size distributions in Figs. 6a and 6b are inferred. Dashed lines: CONTIN analysis of the autocorrelation functions using the MATLAB code *rilt* [9,10].

**Figure S7.**
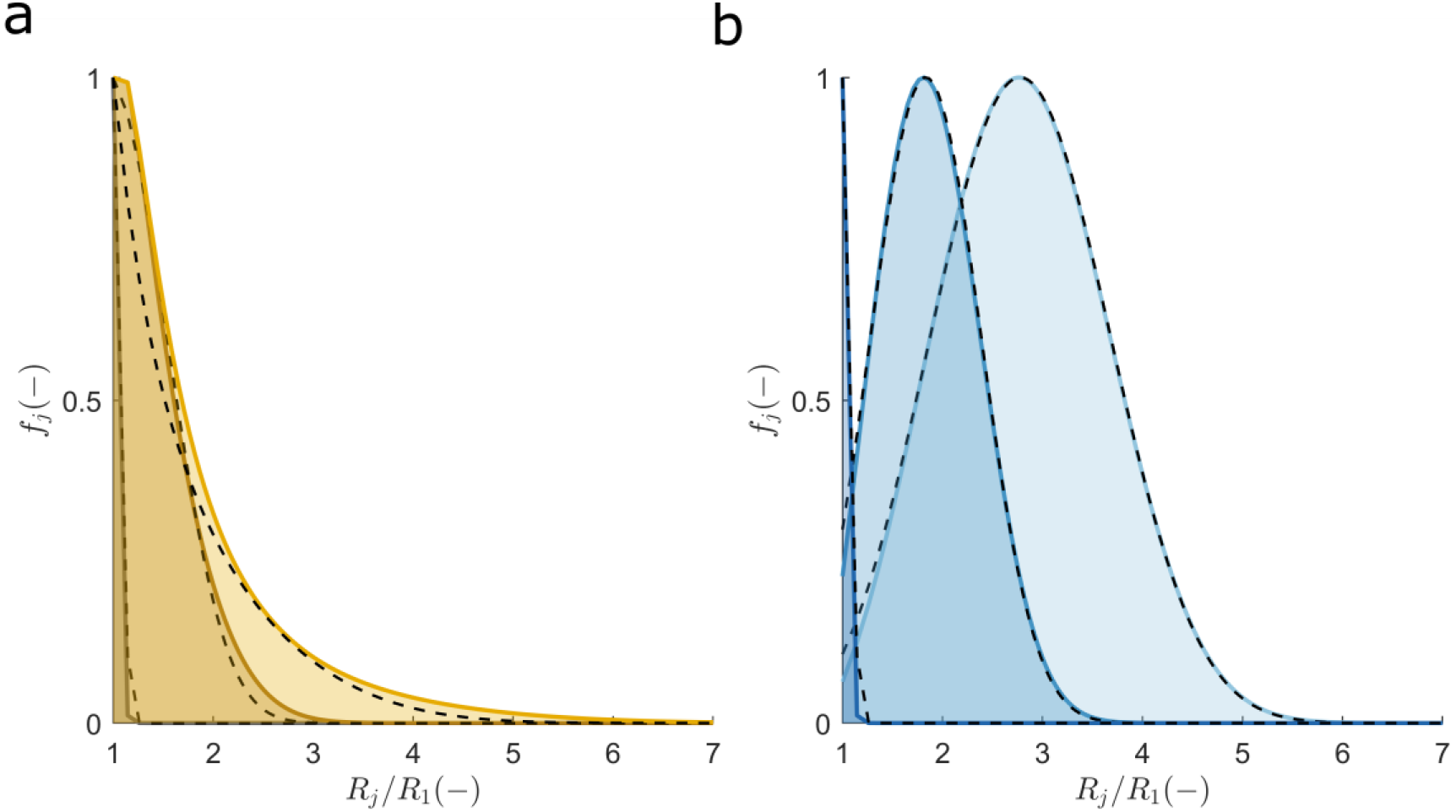
Using the gamma distribution to describe the exact solution of the master model equation (Eq. S2). The endpoint *C_j_*(*t* → ∞) distributions obtained as in (a) Fig. 1d and (b) Fig. S3d are numerically fitted (dashed lines) by the two-parameter gamma distribution and represented in normalized frequency units.

## Notes

### Competing Interest Statement

The authors have declared no competing interest.

